# Explorative Discovery of Gene Signatures and Clinotypes in Glioblastoma Cancer Through GeneTerrain Knowledge Map Representation

**DOI:** 10.1101/2024.04.01.587278

**Authors:** Ehsan Saghapour, Zongliang Yue, Rahul Sharma, Sidharth Kumar, Zhandos Sembay, Christopher D. Willey, Jake Y. Chen

## Abstract

This study introduces the GeneTerrain Knowledge Map Representation (GTKM), a novel method for visualizing gene expression data in cancer research. GTKM leverages protein-protein interactions to graphically display differentially expressed genes (DEGs) on a 2-dimensional contour plot, offering a more nuanced understanding of gene interactions and expression patterns compared to traditional heatmap methods. The research demonstrates GTKM’s utility through four case studies on glioblastoma (GBM) datasets, focusing on survival analysis, subtype identification, IDH1 mutation analysis, and drug sensitivities of different tumor cell lines. Additionally, a prototype website has been developed to showcase these findings, indicating the method’s adaptability for various cancer types. The study reveals that GTKM effectively identifies gene patterns associated with different clinical outcomes in GBM, and its profiles enable the identification of sub-gene signature patterns crucial for predicting survival. The methodology promises significant advancements in precision medicine, providing a powerful tool for understanding complex gene interactions and identifying potential therapeutic targets in cancer treatment.

## Introduction

Understanding the relationship between gene expression and cancer is crucial for the development of innovative diagnostic methods and therapeutic interventions^1,2^. So, gene expression profiling enables the identification of specific gene expression signatures that are indicative of different cancer types, providing a crucial tool for early diagnosis and cancer classification^3,4^. However, analyzing large-scale gene expression datasets can be a challenging task, requiring the application of effective visualization techniques to facilitate data interpretation^5^.

Data visualization is essential for understanding complex datasets, facilitating the identification of patterns, trends, and outliers. In the realm of gene expression analysis, numerous visualization techniques and tools have emerged, such as ClustVis ^6^, MISO ^7^, Mayday ^8^, and GENE-E. These methods encompass straightforward single-figure comparisons of a limited number of features or genes, employing techniques like box plots, bar charts, and heat maps, as well as more advanced techniques like clustering, which group features based on multiple criteria under specific conditions or samples.

Heatmaps have become particularly popular as the preferred method for visualizing gene expression data, often integrating clustering methods that group genes and/or samples together based on the similarity of their gene expression patterns ^5^.

Despite the widespread utilization of heatmaps for visualizing gene associations, existing methods have limitations in inferring relationships based on individual samples. Establishing such relationships is crucial for precision medicine, as it allows for the visualization of associations between multiple genes and a single sample. To address this limitation, we introduce the GeneTerrain Knowledge Map (GTKM) representation – a novel technique designed to capture associations among multiple genes and samples.

The GTKM approach employs Protein-Protein Interactions (PPIs) to graphically depict Differentially Expressed Genes (DEGs) on a two-dimensional (2-D) contour plot. This is achieved by first generating a layout of genes using the DEMA method ^9^, which assigns each gene a unique 2-D location based on its PPIs and pathways. Subsequently, the expression value of each gene is represented by plotting a 2-D Gaussian distribution at its corresponding location in the layout.

In this paper, we present the development and application of the GTKM representation for gene expression analysis. We discuss its advantages over traditional heatmap visualization methods and demonstrate its potential in advancing precision medicine through improved visualization of associations between multiple genes and individual samples.

## Results

Figure 1A illustrates the GeneTerrain method’s framework for this study, which involves generating four GeneTerrains from four samples and a gene expression matrix containing five genes. Subsequently, a protein-protein interaction (PPI) network is constructed and analyzed to establish the genes’ positions in an x-y coordinate system. Ultimately, a two-dimensional Gaussian distribution is mapped based on the gene expression levels and their respective positions. In Figure 1B, the impact of varying sigma values on the GeneTerrain of sample 3 is demonstrated, generating five genes with selected sigma values of 0.05, 0.1, and 0.2.

**Figure 1.**
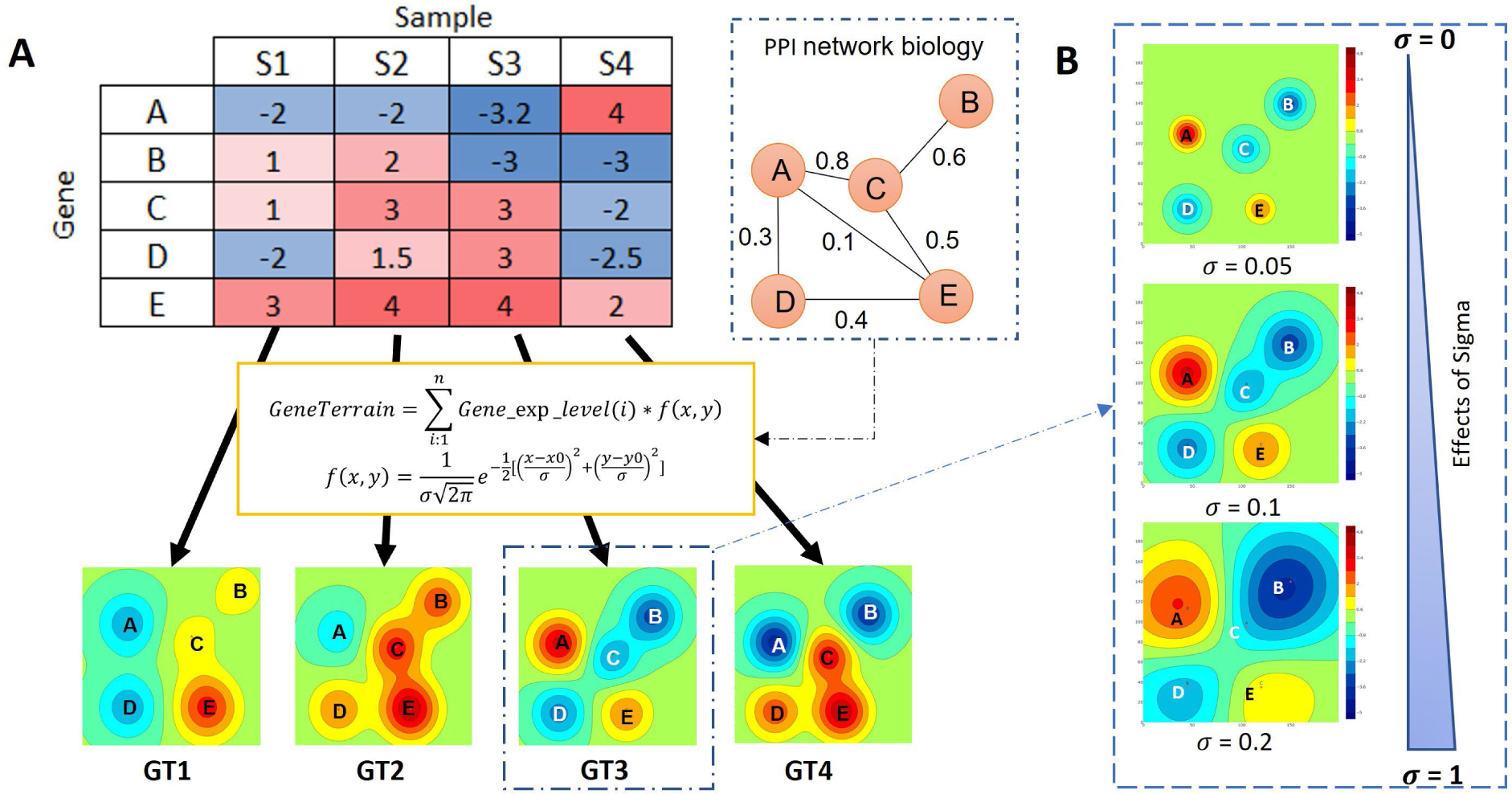
A framework of the Geneterrain method for Cancer. A) Creating four GeneTerrains based on four samples, using a gene expression matrix that includes four samples and five genes. A protein-protein interaction (PPI) network biology is constructed, which will be calculated through PPI to determine the positions of genes in an x-y coordinate system. Finally, a 2D Gaussian distribution will be mapped based on the expression level of the gene and their position. B) Demonstrating the effects of Sigma on GT3, which generates five genes, with selected values of 0.05, 0.1, and 0.2.

### Design GTKM and GTKM profile to discover gene signature

We used three publicly-available GBM datasets obtained from The Cancer Genome Atlas (TCGA)(Dataset ID:HT_HG-U133A)^10^, Chinese Glioma Genome Atlas (CGGA) (Dataset ID: mRNAseq_325) ^11^, and Ivy Glioblastoma Atlas Project (IvyGAP) ^12^.We performed z-score normalization^13^ for all the samples and genes in the data preprocessing. We have conducted three case studies with GBM datasets for (1) Survival analysis, (2) sub-type identification, and (3) IDH1 mutation analysis. Additionally, we have prepared a prototype web app to demonstrate our GBM case studies, which can be replicated for different cancer types.

In these examples, we utilized a list of 2,290 differentially expressed genes (DEGs) for our analysis. These DEGs were identified as differentially expressed between tumor and normal samples, exhibiting a log2 fold-change of ≥ 2 and a false discovery rate (FDR) of ≤ 1.0e-3, as obtained from OncoDB ^14^. We subsequently extracted protein-protein interactions (PPIs) from the DEG list, employing a confidence cutoff of 0.9 at the highest PPI quality level. This process yielded a total of 4,612 PPIs involving 1,163 candidate genes. To maintain network integrity, we excluded small networks not connected to the primary (largest) network, resulting in a final set of 912 candidate genes. The DEMA algorithm ^9^ was then employed to generate a network layout using these candidate genes. Ultimately, we applied the default coefficient ratios in DEMA as kb/ka= 912 and Kc = 9,120 for the DEMA layout (Figure 1 in Supplementary 1).

We generated GTKMs and GTKM Profiles for three case studies to identify conserved patterns of gene regulation (Figure 2 in Supplementary 2). The GTKMs were visualized using varying shades of blue and red, with the darkest blue and red fields representing the highest downregulated and upregulated intensities, respectively. The signal background of the GTKMs was denoted by different shades of green. To effectively illustrate high similarity signal areas, we employed a technique called Levelized colors to discretize the GTKMs, allowing for effortless identification and quantification of regions of interest. Additionally, we plotted contour lines using predefined shades of blue, red, and green colors to enhance the GTKMs’ visual representation. The GTKMs revealed three regions: upregulated, downregulated, and neutral. Upregulated and downregulated regions signify the convergence of gene expression in a local area, while the neutral region indicates an absence of convergence. This information aids in pinpointing key regions where specific properties prevail.

**Figure 2.**
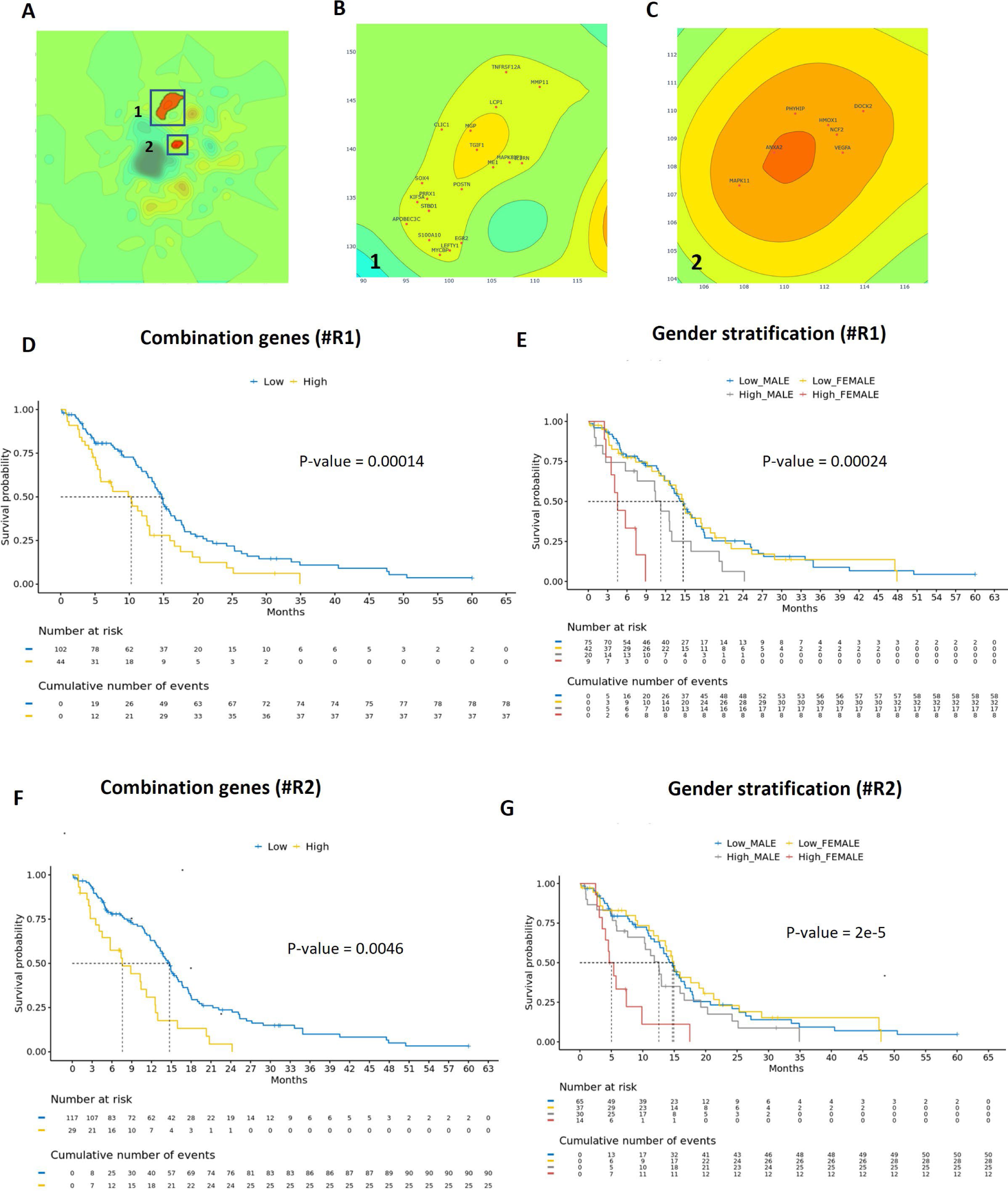
Flowchart of GTKM profiles of case study 1. A) Displaying the contour of the GTKM profile for short survival as the background and identifying two regions in upregulated areas with significant p-values based on log-rank. B) Showing region 1 in the contour of the short survival GTKM profile and the names of their genes in upregulated areas. C) Showing region 2 in the contour of the short survival GTKM profile and the names of their genes in upregulated areas. D) Kaplan-Meier curve plot based on the combination of genes from region 1 to stratify two groups by low and high gene expression levels. E) Kaplan-Meier curve plot based on the combination of genes from region 1 to stratify gender by low and high gene expression levels. F) Kaplan-Meier curve plot based on the combination of genes from region 2 to stratify two groups by low and high gene expression levels. G) Kaplan-Meier curve plot based on the combination of genes from region 2 to stratify gender by low and high gene expression levels

GTKMs uncover gene patterns that facilitate the classification of glioblastoma multiforme (GBM) cohorts with poor survival, GBM molecular subtypes, and IDH1 mutation. Specifically, three case studies were conducted to extract clinotype-related gene patterns using the GTKM exploration tool. In the first case study, an investigation was carried out to examine gene signature patterns pertinent to survival prediction. In the second case study, an exploration was conducted to identify gene signature patterns associated with GBM molecular subtypes. In the third case study, gene signature patterns in relation to IDH1 mutation were investigated. The GTKMs serve as a powerful tool for visualizing and analyzing complex data sets, ultimately leading to the recognition of key gene patterns associated with specific clinical outcomes.

### GTKM profile identifies sub-gene signature patterns that improve predicting survival

The gene patterns related to survival, as extracted from GTKMs, indicate disrupted molecular mechanisms in patients with poor survival outcomes. The construction of survival-related GTKM profiles incorporates survival information. Patients were divided into short and long survival groups based on the average survival duration for glioblastoma patients (8 months or 240 days), as per the National Brain Tumor Society. The short survival group comprised 176 patients, while the long survival group consisted of 348 patients. Figure 1 in Supplementary 2 displays GTKM profiles and their contours for both long and short survival patients. The color scale used ranges from −1 to 1, representing continuous values for the GTKMs. A step of 0.1 has been set between each contour level. Three distinct regions can be observed in the GTKMs: upregulated, downregulated, and neutral regions. The log-rank test p-value, calculated based on the intensity of each region between the two groups, uncovers survival-related gene patterns. Two upregulated regions (1 and 2) are identified, and these regions are derived from the short survival contour GTKM (Figure 2A).

Region 1 and region 2 encompass 19 genes and 6 genes, respectively (Figure 2B-C). Subsequently, cSurvival is employed, an advanced framework for studying biomarker interactions in cancer that includes a user-adjustable pipeline, a curated and integrated database, and several key features. One of these features is the capability to perform joint analysis with two genomic predictors to identify interacting biomarkers, utilizing new algorithms to determine optimal cutoffs for these predictors. Another feature is the capacity to conduct survival analysis at both the gene and gene set (GS) levels. In region 1, the TNFRSF12A gene exhibited the most significant p-value (log-rank p = 0.0029) in the Kaplan-Meier plot, while the PRRX1 gene had the least significant p-value (log-rank p = 0.53) (Figures 2-20 in Supplementary 2). Figure 2D presents a Kaplan-Meier curve based on combinations of genes in region 1, with a p-value of 0.0042. High expression levels of these genes are associated with poor overall survival. Moreover, the analysis indicated that patients with lower expression levels of these genes have a 50% probability of a survival rate for 15 months, while those with higher expression levels had a probability of a survival rate for only 10 months. Yu Zhang et al.^15^ made a similar discovery, noting that TNFRSF12A expression levels were elevated in recurrent and secondary gliomas compared to primary gliomas. Additionally, they found that TNFRSF12A exhibited higher expression in wildtype and 1p19q non-coding gliomas when compared to IDH mutants or 1p19q co-deletions.

In region 2, the MAPK11 gene exhibited the most significant p-value (log-rank p = 0.037) in the Kaplan-Meier plot, while the HMOX1 gene had the least significant p-value (log-rank p = 0.25) (Figures 21-26 in Supplementary 2). When all of these genes were combined, the overall p-value became much more significant (log-rank p = 0.00014), as depicted in Figure 2F. High expression of these genes is associated with poor overall survival in GBM patients. Furthermore, the analysis demonstrates that patients with lower expression levels of genes have a 50% survival probability of 14 months, compared to patients with higher expression levels of these genes who have a survival probability of only 8 months. Recently, D.M. Fernandez-Aroca et al. ^16^ conducted research revealing that MAPK11 plays a pivotal role in the cellular response to ionizing radiation. Their findings suggest that MAPK11 is involved in controlling senescence and may have implications for cancer cell survival.

### Gender related genes associated with worse female survival rate

We plotted Kaplan-Meier survival curves for both female and male patients based on the expression levels of genes in two regions. The results show that female patients with high expression levels of these genes have a poor prognosis, with a survival time of only 5 months (50% survival probability), compared to patients with low expression levels of these genes who have a survival time of 15 months (50% survival probability) (Figure 2E-G).

Additionally, we expanded the number of genes to 27 through the DEMA layout in region 1 (Figure 27 in Supplementary 2) and generated a local GTKM profile of short survival time (sigma = 0.01). By using the local GTKM profile, we can expand the number of genes and explore gene-gene interactions.

### GTKM profile shows unique patterns for each cancer subtype

We constructed four GTKM profiles for the previously described GBM molecular subtypes, including Proneural, Mesenchymal, Classical, and Neural (believed to be artificial due to normal brain tissue contamination, i.e., a highly invasive tumor) subtypes, using three datasets: TCGA, CGGA, and IvyGAP. These profiles displayed significant distinct molecular functional indications. The TCGA and CGGA datasets included microarray gene expression, while IvyGAP featured RNA-Seq gene expression. Our aim is to demonstrate the effectiveness of this approach for different types of datasets (Figure 3A).

**Figure 3.**
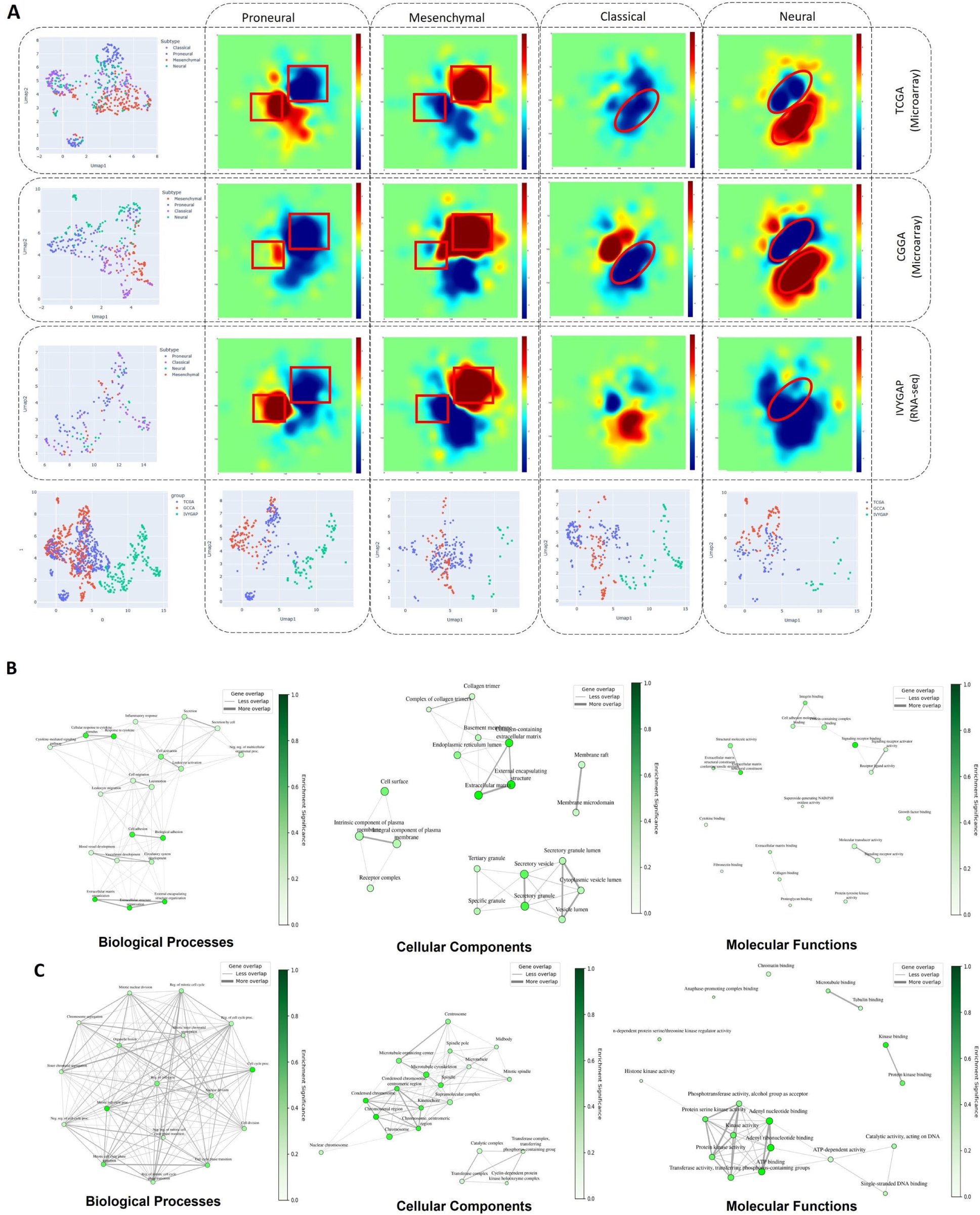
Flowchart of GTKM profiles of case study 2. A) GTKM profiles of three datasets, including TCGA, CGGA, and IvyGAP, for each cancer subtype in TCGA-GBM and CGGA datasets. B) The network-based pathway enrichment analysis, including gene ontology (GO) for biological processes, cellular components, and molecular functions for the gene list in the upregulated and downregulated areas in Mesenchymal and Proneural GTKM profiles, respectively. C) The network-based pathway enrichment analysis, including gene ontology (GO) for biological processes, cellular components, and molecular functions for the gene list in the downregulated and upregulated areas in Mesenchymal and Proneural GTKM profiles.

Moreover, Figure 3A reveals that the Proneural and Mesenchymal GTKM profiles for the three datasets are similar in two areas, which are highlighted by red squares. These similarities can be considered as unique patterns for these subtypes and their related genes. Another significant observation is that the roles of genes in the two subgroups are opposite to each other. An upregulated region has been identified in the GTKM profiles for the Classical subtype based on the TCGA and CGGA datasets, while the Classical GTKM profile of IvyGAP differs. In the Neural subtype, an upregulated region has been identified in the GTKM profiles for the Neural subtype in all three datasets, with an upregulated area based on the GTKM profiles of the CGGA and TCGA datasets. To demonstrate the effectiveness of this approach, we applied the UMAP method to each dataset to classify the subtypes. Additionally, we mapped the results to each subtype and plotted them.

To extract pathways for the Proneural and Mesenchymal subtypes, we compiled a list of gene names in two interval values—the highest upregulated and lowest downregulated areas. This approach was based on their GTKM profiles for the three datasets, which showed that these regions are associated with the desired subtypes. Figure 3B illustrates the network-based pathway enrichment analysis, including the gene ontology (GO) biological process, cellular component, and molecular function for the gene list in the upregulated area in Mesenchymal and the downregulated area in Proneural GTKM, respectively. Figure 3C displays the network-based pathway enrichment analysis, including the gene ontology (GO) biological process, cellular component, and molecular function for the gene list in the downregulated area in Mesenchymal and upregulated area in Proneural GTKM profiles. Figures 3B and 3C were generated using the ShinyGO (V0.7) application^17^.

For building GTKM profiles of TCGA, there are 142, 156, 138, and 87 samples for Classical, Mesenchymal, Proneural, and Neural subtypes of TCGA-GBM cancer, respectively. For building GTKM profiles of CGGA, there are 73, 68, 100, and 80 samples for Classical, Mesenchymal, Proneural, and Neural subtypes, respectively. For building GTKM profiles of IvyGAP, there are 56, 17, 67, and 17 samples for Classical, Mesenchymal, Proneural, and Neural subtypes, respectively. Additionally, since some samples in the IvyGAP dataset have two labels, we have removed them. The values of the color scale of GTKM profiles are considered between −5 and 5.

Our study has revealed that extracellular matrix organization, external encapsulating structure, and signaling receptor binding are crucial pathways in the regulation of the Mesenchymal subtype in the upregulated area and the Proneural subtype in the downregulated area (Supplementary 3). It is noteworthy that in the biological process category, the “cytokine response” and “cytokine-mediated signaling pathway”, in the cellular component category, the “external encapsulating structure” and in the molecular function category, “signaling receptor binding” have all been activated in mesenchymal cells. This observation strongly supports the idea that mesenchymal cells serve as cytokine carriers^18^. Significantly, cytokine signaling plays a crucial role in the transition to cancer cells through the recruitment of mesenchymal stem cells (MSCs). Furthermore, MSCs facilitate the migration, invasion, and adhesion of cancer cells. Additionally, cell cycle processes, chromosomal regions, and ATP binding are identified as key pathways in the regulation of the Proneural subtype in the upregulated area and the Mesenchymal subtype in the downregulated area. Proneural cells exhibit distinctive cytoskeletal functions during subtype transitions to mesenchymal cells in GBM ^19^. These proneural cells display heightened activities in cell cycle and cell division within biological processes, microtubule cytoskeleton involvement within cellular components, and ATP-dependent activity for energy supply, implying significant morphological changes. These pathways were determined by analyzing enrichment FDR values for GO-BP, GO-CC, and GO-MF categories, respectively (Supplementary 4).

### GTKM profiles identifies sub-gene signature patterns based on gender-specific IDH1 mutation

Isocitrate dehydrogenase (IDH) is a key rate-limiting enzyme that participates in several major metabolic processes, such as the Krebs cycle, glutamine metabolism, lipogenesis, and redox regulation. IDH mutations are directly associated with the onset and progression of glioma, making it a significant prospective therapeutic target ^20^. The two sample groups, IDH1-wildtype and IDH1-mutant (R132G, R132C, R132H), are based on IDH1 mutation and include 370 wildtype samples and 30 IDH1-mutated samples from the TCGA dataset. The GTKMs display significantly different gene patterns (Figure 4A). The GTKM of IDH1-mutant consists of one upregulated area and one downregulated area, while the GTKM of IDH1 wildtype includes one upregulated area. The scale of the GTKMs ranges between −3 and +3. GTKMs display distinct gene expression profiles for cases belonging to upregulated vs. downregulated wildtype/mutant groups. For comparison based on the GTKM of wildtype, 365 genes were identified with a threshold value greater than 1 in the upregulated area. Similarly, for the upregulated and downregulated areas of the GTKM of mutant cases, 353 and 343 genes were identified, respectively. Additionally, we constructed gender-specific GTKM profiles for IDH1-Mutant patients to explore potentially significant genes. We identified a region of 92 genes that were upregulated and downregulated simultaneously based on the overlap of the two GTKM profiles (Figure 4B). Out of the 92 genes, 31 were significantly associated with survival using cSurvival analysis ^21^. Indeed, the p-values of NCAPH, AURKB, TOP2A, and CDK2 genes were less than 0.00001. We then extracted the top five KEGG pathways associated with these genes using gProfiler. Additionally, we calculated the mean of gene expression values for both groups. Further analysis revealed that out of the 31 significant genes, 28 showed that females with low gene expression had a poorer prognosis compared to other categories (Figure 4C). Interestingly, the four genes highly associated with survival are also critical components of the Chromosome Condensation Pathway. This pathway elucidates the roles of AURKB, TOP2A, CDK2, and NCAPH in chromosome condensation, a ubiquitous process in most eukaryotic cells^22^. Chromatin condensation during mitosis, with a specific focus on these major molecular players, initiates and maintains the particular chromatin conformation. Furthermore, evidence suggests that interference with chromatin condensation leads to an inadequate activation of the DNA damage response (DDR), ultimately reducing cell recovery and survival^23^.

**Figure 4.**
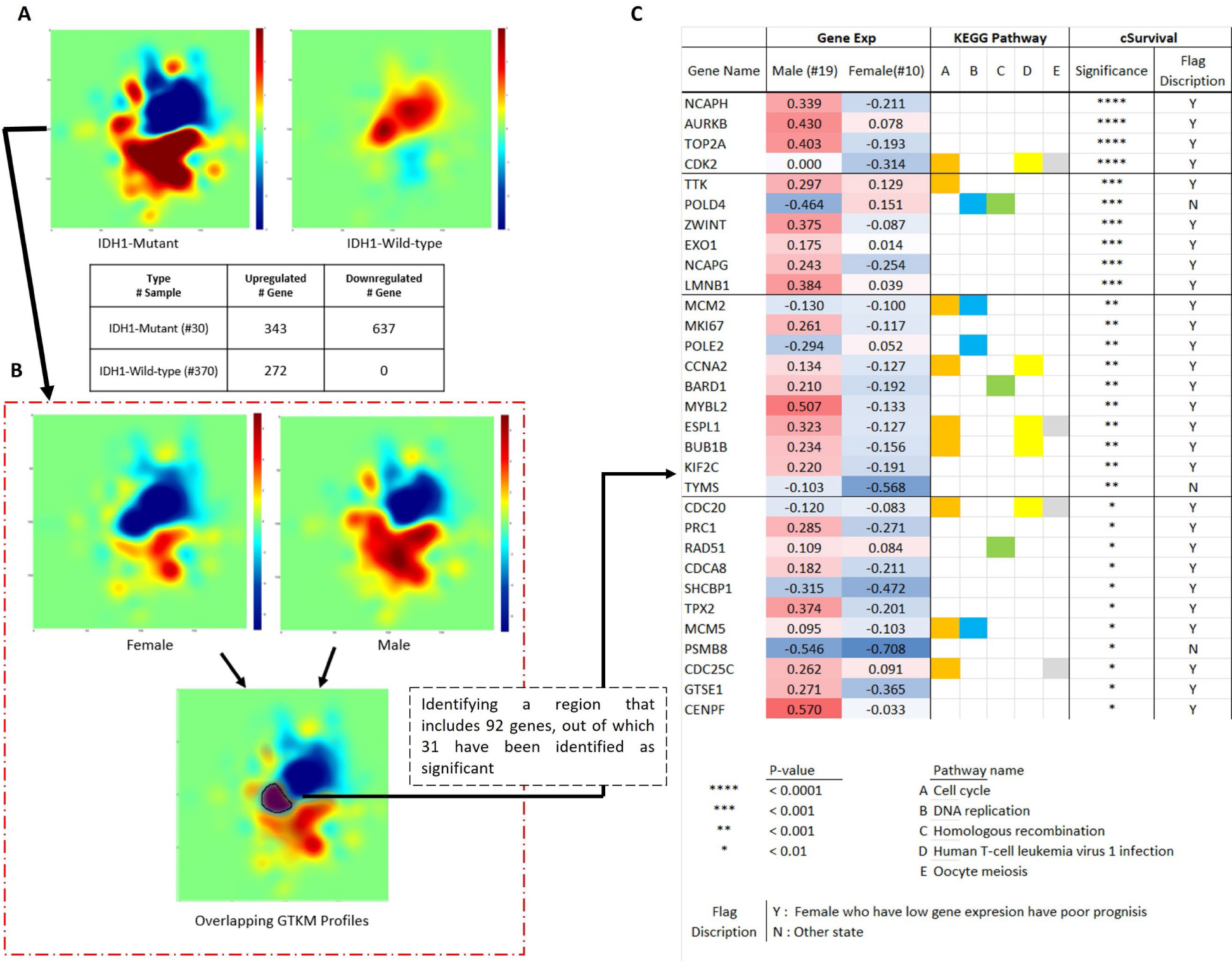
Flowchart of GTKM profiles of case study 3. A) Constructing GTKM profiles for IDH1-mutant and IDH1 wild-type samples, as well as displaying the number of genes in upregulated and downregulated areas within the GTKM profiles. B) Constructing GTKM profiles for male and female IDH1-mutant samples, and identifying a region containing 92 genes that play a role in both upregulated and downregulated areas simultaneously in IDH1-mutant samples based on gender by overlapping male and female GTKM profiles. C) Displaying 31 out of 92 genes with significant p-values based on gender stratification using cSurvival, as well as the mean gene expression values for both male and female groups in IDH1-mutant samples.

### GTKM profiles identifies sub-gene signature patterns based on drug sensitivities of different tumor cell lines

Drug resistance is a major obstacle in cancer treatment, diminishing the effectiveness of chemotherapy and targeted therapies. The molecular mechanisms driving drug resistance evolution remain unclear, requiring sophisticated methods for exploration. In our case study, we analyzed gene expression dynamics in tumor cells treated with db-cAMP, aiming to uncover mechanisms of drug resistance in cancer. We analyzed three glioblastoma cell lines (DBTRG-05MG, U87, LN18) treated with db-cAMP, using gene expression data from the GSE128722 dataset over 0-48 hours. For this case study, we selected 1000 highly variable genes, from which 903 were selected for network analysis based on their correlation profiles with threshold value of 0.9. Then, these genes were used to construct a gene interaction network via a force-directed algorithm method. Then GTKM profiles were then generated for each tumor cell line across the specified time intervals to visualize the gene expression landscapes. The GeneTerrain Knowledge Maps highlighted the dynamic gene expression changes due to dbcAMP treatment. The DBTRG-05MG cell line showed significant temporal gene expression shifts, while U87 and LN18 displayed minimal changes, indicating possible intrinsic resistance. Identifying key gene clusters with significant expression changes is crucial for understanding the mechanisms of drug resistance and identifying potential biomarkers for treatment sensitivity. We identified 18 genes with altered expression in the D5 time point for DBTRG-05MG, including “S100A6”, “GDF15”, “RPL37A”, “TIMP1”, “SPARC”, “ATOX1”, “HSPA5”, “BNIP3”, “CDC20”, “H2AFX” “HSPA9”, “TCEB2”, “PTBP1”, “NDUFS8”, “NDUFB9”, “ERGIC3”, “RPLP1”, and “TECR”. These genes exhibited changes in gene expression within the two boxed areas of interest. Also, 6 genes including “FTL” “COX5B” “EZR” “RSAD2” “ISOC2” and “LDHB” are identified for D5 time point for U87.

Figure 5 presents comparative GTKMs to visualize the gene expression dynamics across three tumor cell lines—DBTRG-05MG, U87, and LN18—following treatment with 1 mol/L db-cAMP. Observations are mapped at 0, 6, 12, 24, and 48 hours post-treatment. Key details include: Each row corresponds to a different cell line. Time points for DBTRG-05MG are labeled D1 to D5, for U87 as U1 to U5, and for LN18 as L1 to L5. DBTRG-05MG shows notable temporal shifts in gene expression. U87 and LN18 exhibit distinct gene expression patterns, reflecting their unique responses to the treatment. Boxed sections highlight significant gene activity areas, which could be associated with drug resistance mechanisms.

**Figure 5.**
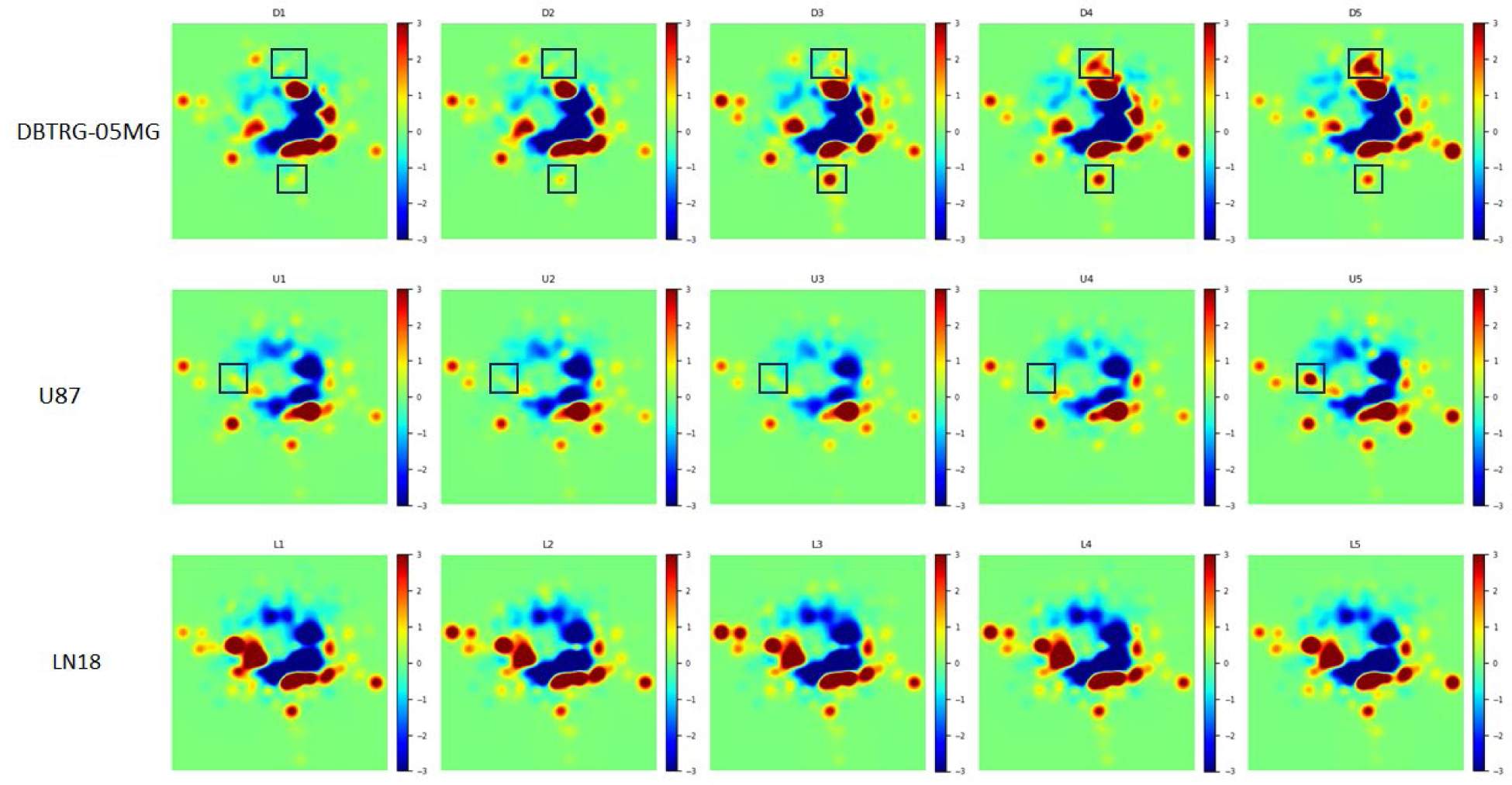
Flowchart of GTKMs of case study 4. Comparative GTKMs illustrating the gene expression dynamics in DBTRG-05MG, U87, and LN18 tumor cell lines over time intervals of 0, 6, 12, 24, and 48-hours post-treatment with 1 mol/L db-cAMP. Each row represents a different cell line with sequential time points labeled from D1 to D5 for DBTRG-05MG, and U1 to U5 for U87, and L1 to L5 for LN18 respectively. Color gradients indicate levels of gene expression, with red representing upregulation, blue downregulation, and green minimal to no change. Notable temporal shifts are highlighted in the DBTRG-05MG cell line, whereas U87 and LN18 demonstrate distinct patterns of gene expression, reflecting their unique adaptive responses to the drug treatment. The boxed areas emphasize regions with significant gene activity potentially linked to drug resistance mechanisms.

For example, this high expression of S100A6 in Cancer stem cells (CSCs) is significant because CSCs are known for their self-renewal, pluripotency, tumor initiation, and resistance to radiotherapy or chemotherapy, which can be the cause of cancer recurrence. (https://biomarkerres.biomedcentral.com/articles/10.1186/s40364-023-00515-3)

GDF15 has been identified as playing a crucial role in promoting a stem cell-like phenotype in glioma stem cell-like cells (GSCLCs). This is achieved through the regulation of ERK1/2–c-Fos– LIF signaling, which enhances the expression of glioma stem cell (GSC) markers and increases the sphere-formation capability and cell division in GSCLCs. (https://pubmed.ncbi.nlm.nih.gov/34697897/) (https://www.nature.com/articles/s41420-020-00395-8)

## Discussion

Visualization is one of the most common approaches for analyzing biological data. Clustering, dimension reduction, and heatmaps are prominent methods used to analyze systems biology data. Among these three methods, only heatmaps utilize multiple genes and samples to capture biological processes for making inferences, such as survival analysis. However, heatmaps have two limitations: (1) they are not meaningful with a single patient sample, and (2) they rely on clustering methods to map the associations among genes (or proteins) as biological relationships, rather than using network biology. This article proposes a novel methodology called GeneTerrain Knowledge Map representation (GTKM), which can plot visualizations for single and multiple patient samples while establishing relationships among genes (or proteins) using network biology. GTKM is similar to a 2D contour plot, where every gene (or protein) has a fixed location (coordinate) on the map. These gene (or protein) positions (or layouts) are derived from protein-protein interaction-based network biology and force-directed algorithms. In this article, we used DEMA to construct the layout of gene (or protein) positions on the 2D plot. GTKM offers flexibility in selecting algorithms to create the layout; thus, instead of DEMA, we could also use layouts such as Kamada-Kawai ^24^, Bayesian ^25^, Spring ^26,27^, and Spectral ^28^.

Through four case studies using GBM data, we demonstrate that the GeneTerrains produced using GTKM are highly beneficial for: (1) survival analysis, (2) cancer subtype identification, and (3) specific gene-based (IDH1) mutation analysis, and 4) drug sensitivity to extract informative genes and pathways. For GBM survival analysis, the results highlight two aspects: 1) Consideration of Multiple Genes: The study emphasizes the importance of considering multiple genes simultaneously in survival analysis for GBM. This approach enables the identification of gene patterns related to survival outcomes, which can help develop personalized treatment strategies for GBM patients based on their unique gene expression profiles. By analyzing multiple genes, researchers can gain a better understanding of the genetic factors that influence survival rates in GBM patients, potentially leading to more effective treatment options. 2) Gender-Related Genes: We demonstrated that gender-related genes play a crucial role in determining the survival rates of GBM patients. By identifying specific genes and interactions involved, researchers can gain valuable insights for developing personalized treatment strategies. Understanding the genetic factors that influence survival rates can help clinicians tailor treatment plans to individual patients, potentially leading to better outcomes. These findings may also have broader implications for understanding the role of gender in the development and progression of GBM.

For cancer subtype identification, this study provides valuable insights into the genomic landscape of GBM cancer and demonstrates the effectiveness of using multiple datasets to identify unique patterns in different subtypes. We found that the Proneural and Mesenchymal GTKM profiles were similar in two areas across all three datasets. These similarities can be considered as unique patterns for these subtypes and their related genes.

For IDH1 mutant and wild-type analysis, we demonstrated that the gene expression patterns of IDH1-wildtype and IDH1-mutant groups are significantly different, with the mutant group encompassing both upregulated and downregulated areas. Additionally, we constructed gender-specific profiles for IDH1-mutant patients and identified 92 genes by overlapping male and female GTKM profiles. Out of these, 28 were significant and associated with a poorer prognosis for female patients who had high gene expression values based on the RNA-seq of TCGA-GBM dataset.

For drug sensitivities of different tumor cell lines, underscore the importance of GTKMs in understanding gene expression dynamics across tumor cell lines in response to drug treatment. The distinct patterns observed in each cell line, particularly the notable shifts in DBTRG-05MG as opposed to the minimal changes in U87 and LN18, highlight the utility of this approach in deciphering the complex molecular landscape of cancer cells and their varied responses to chemotherapy. These insights pave the way for more targeted and effective therapeutic strategies, tailored to the specific genetic makeup of different tumor types.

Overall, GTKM is an innovative method, and we believe it can be employed in the unsupervised and visual analysis of various types of cancer and other diseases. This method can be utilized to visualize high-dimensional data and identify unique patterns or clusters within the data. Additionally, GTKM can plot gene expressions of individual patients, showcasing its potential for precision medicine applications. In the future, we intend to use GeneTerrain to infer cancer progression based on patients’ pre- and post-treatment gene expression.

## Acknowledgments

We would like to thank Thanh Nguyen for his valuable suggestions regarding the case studies in the paper. The work was in part supported by the internal University of Alabama at Birmingham research grants to JC, the National Institutes of Health grant awards U54TR001005 in which JC serves as a co-investigator, and R01 awards R01HL150078 in which RW serves as principle investigator and JC serves as co-investigator.

## Data availability

All datasets used in this study are freely accessible from public sources. Cancer Genome Atlas (TCGA): https://portal.gdc.cancer.gov/, Chinese Glioma Genome Atlas (CGGA): http://www.cgga.org.cn/, Ivy Glioblastoma Atlas Project (IvyGAP): http://glioblastoma.alleninstitute.org/, GSE128722: Available through the Gene Expression Omnibus (GEO) at https://www.ncbi.nlm.nih.gov/geo/query/acc.cgi?acc=GSE128722

## Code availability

The code for implementing the proposed methodology is available at https://github.com/aimed-lab/GTKM

## Materials and Methods

### The Workflow in GTKM generation

We developed a three-step procedure for performing visual analytics on GBM microarray datasets. The input for this procedure is a list of differentially expressed genes (DEGs) from a specific biological condition. First, we extracted protein-protein interactions (PPIs) from HAPPI 2.0 ^28^ and the BEERE web server ^29^ to construct a weighted gene network. Second, we generated a network layout with quantitatively measured biological features from the weighted gene network using the DEMA algorithm ^9^. Third, we applied the GeneTerrain technique ^30^ to form GTKMs by featuring gene signature extraction with signal interpolation using a 2D Gaussian distribution and signal denoising using contour lines.

### Network layout generation using DEMA

To provide an unbiased network layout that characterizes and optimizes GBM gene modules with high modularity and clarity, we adopted the distance-bounded energy-field minimization algorithm (DEMA). In the GBM’s DEMA layout, we utilized two attributes: (1) the retrieved 5-star Protein-Protein Interactions with H-scores ≥ 0.9 from HAPPI 2.0 (5) and the BEERE web server (6) were used as interaction coefficients in the edge-weighted property, and (2) the enriched pathways with P-values ≤ 0.05 from PAGER ^31^ using the candidate gene list. We executed the DEMA layout generation via Cytoscape software ^32^.

### Visual Exploration using GeneTerrain

The GeneTerrain technique, introduced by You et al. ^30^, was applied to generate GTKMs for forming biological hypotheses. The principle of GeneTerrain’s signal interpolation is shown in Figure 3A in Supplementary 1. There are two primary inputs for GeneTerrain: (1) gene signal, and (2) gene network layout (graph). We imported the normalized gene expression as the gene signal input. After applying network generation using DEMA, we extracted the gene coordinates as layout input. Upon executing GeneTerrain, a 2-D graph is yielded, in which each point represents a gene and a color field where the gene signal interpolates using a 2D normal Gaussian distribution (or gene representation).

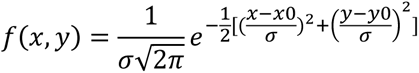

Where, *x*_0_ and y_0_ define the coordinates of a gene, and Sigma (*σ*) is a customized parameter ranging from 0 to 1. Sigma plays an essential role in visually demonstrating the effect on space. The larger the Sigma, the more significant the node’s effect in 2D space. The pixel signal is calculated based on the sum effect of nodes using 2D Gaussian distribution.

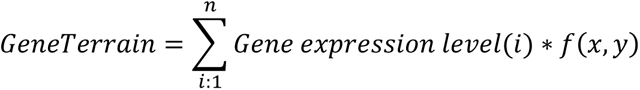

Figure 3A in Supplementary 1 shows the scale of Sigma and its effect on the surface signal in the demo. The layout is generated by DEMA using a sample from the GBM dataset.

In a GTKM, we applied the “rainbow” color palette to represent positive, negative, or zero synergetic signals in red, blue, or green, respectively. If the synergetic signal of a geneset (or gene) is 0, the color field will appear green in GTKM. Moreover, if there is no divergence between gene signals, the region will also appear green. GTKM is a GeneTerrain built using the gene expressions of an individual patient. The patient-specific GTKMs are further superimposed on one another to create a GTKM Profile based on the pixel-level average (a unified GTKM for a group of patients), as shown in Figure 2. We applied the same color conventions to GTKM Profile.

