## supplementary for "Explorative Discovery of Gene Signatures and Clinotypes in Glioblastoma Cancer Through GeneTerrain Knowledge Map Representation": Supplementary 1.pptx

### Slide 1
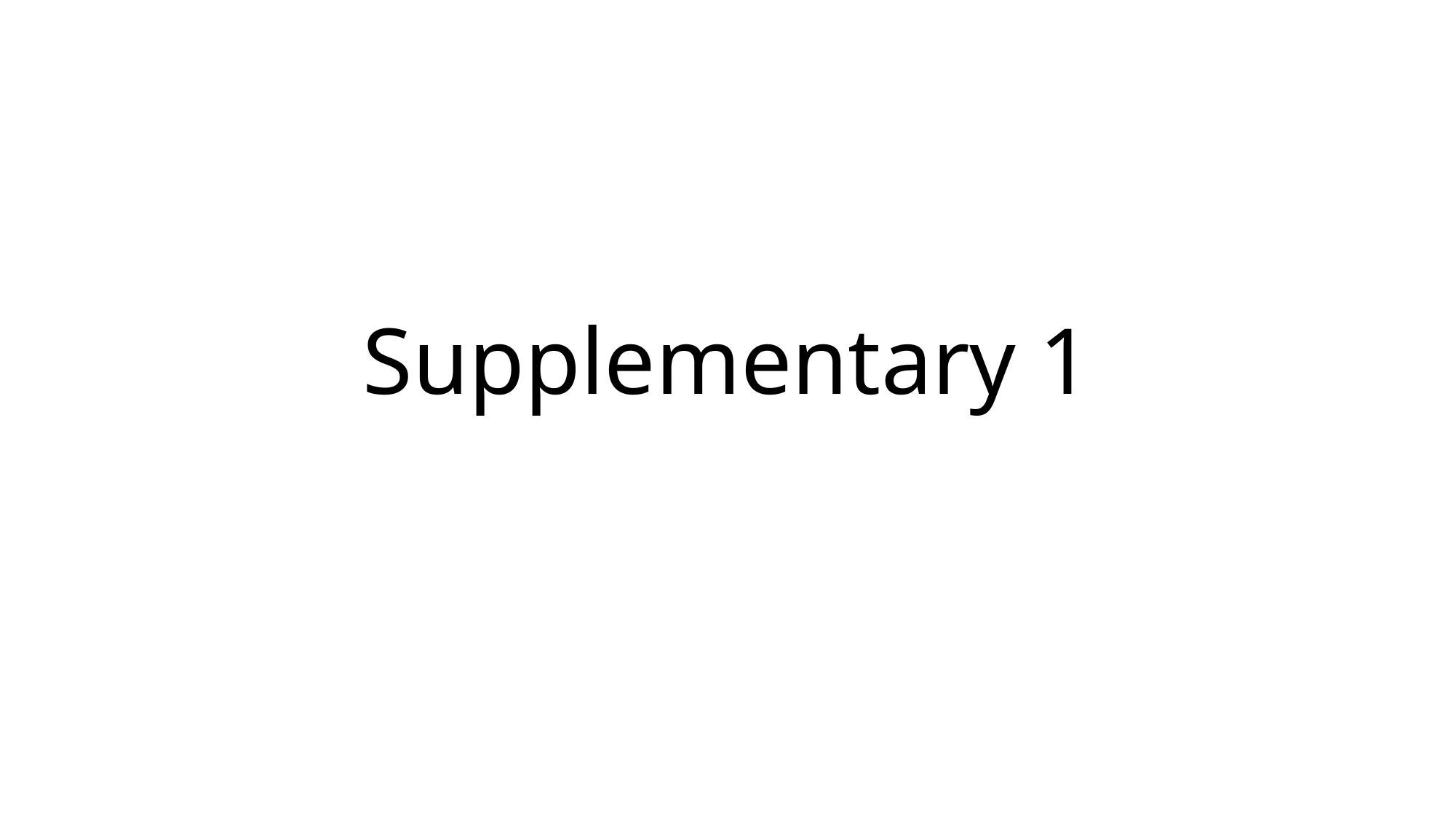

#### Supplementary 1

### Slide 2
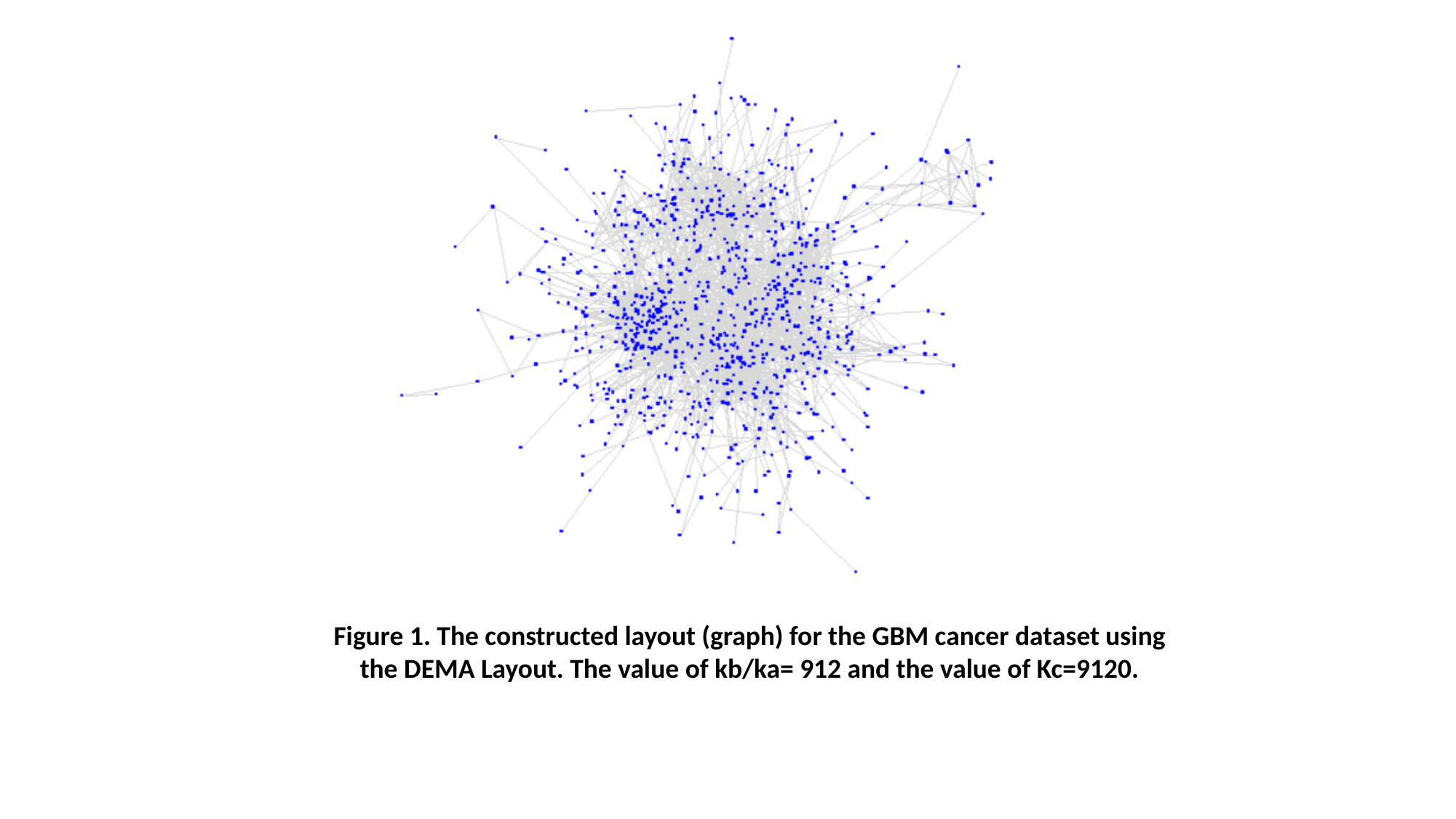

Figure 1. The constructed layout (graph) for the GBM cancer dataset using the DEMA Layout. The value of kb/ka= 912 and the value of Kc=9120.

### Slide 3
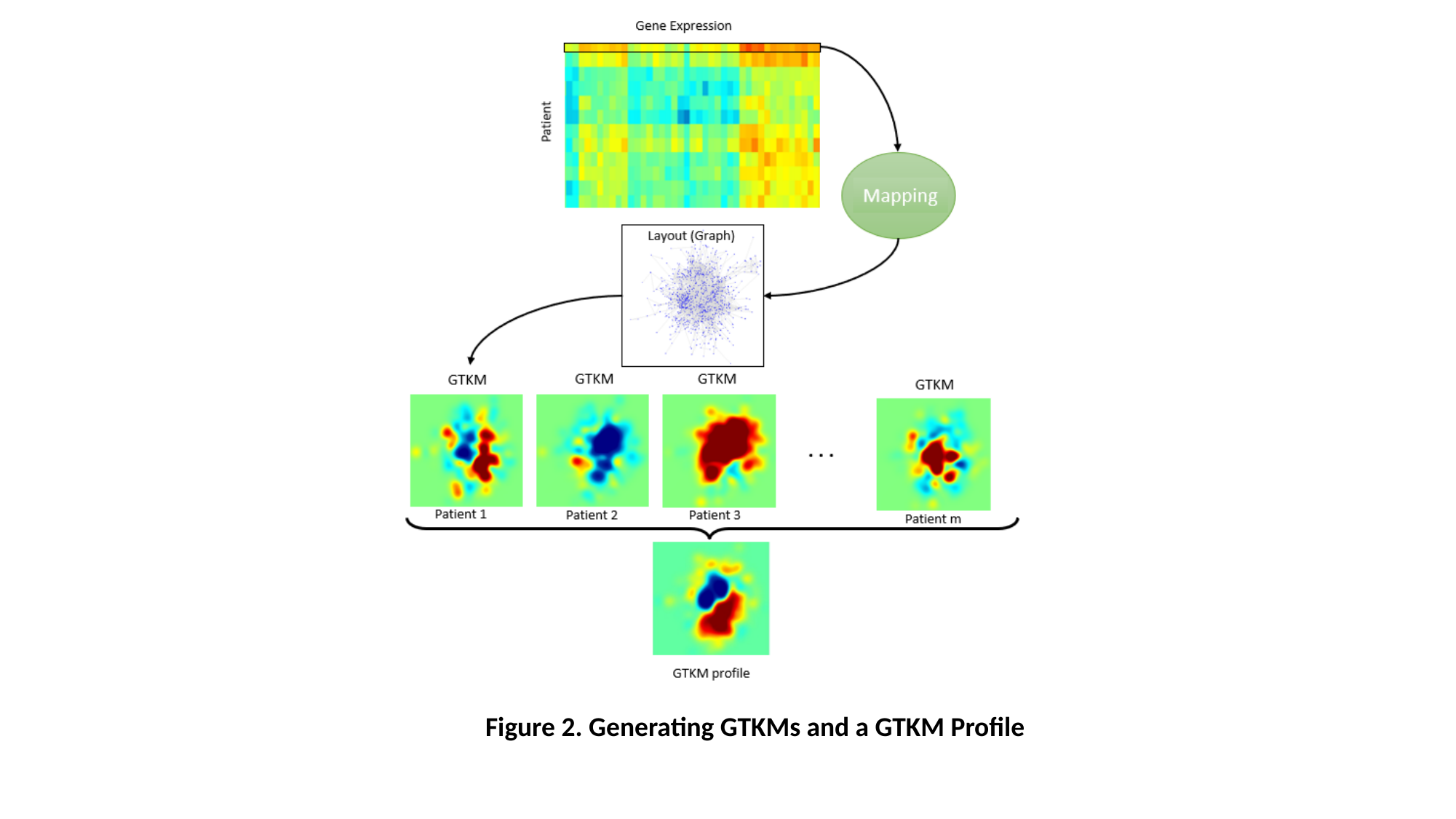

Figure 2. Generating GTKMs and a GTKM Profile

### Slide 4
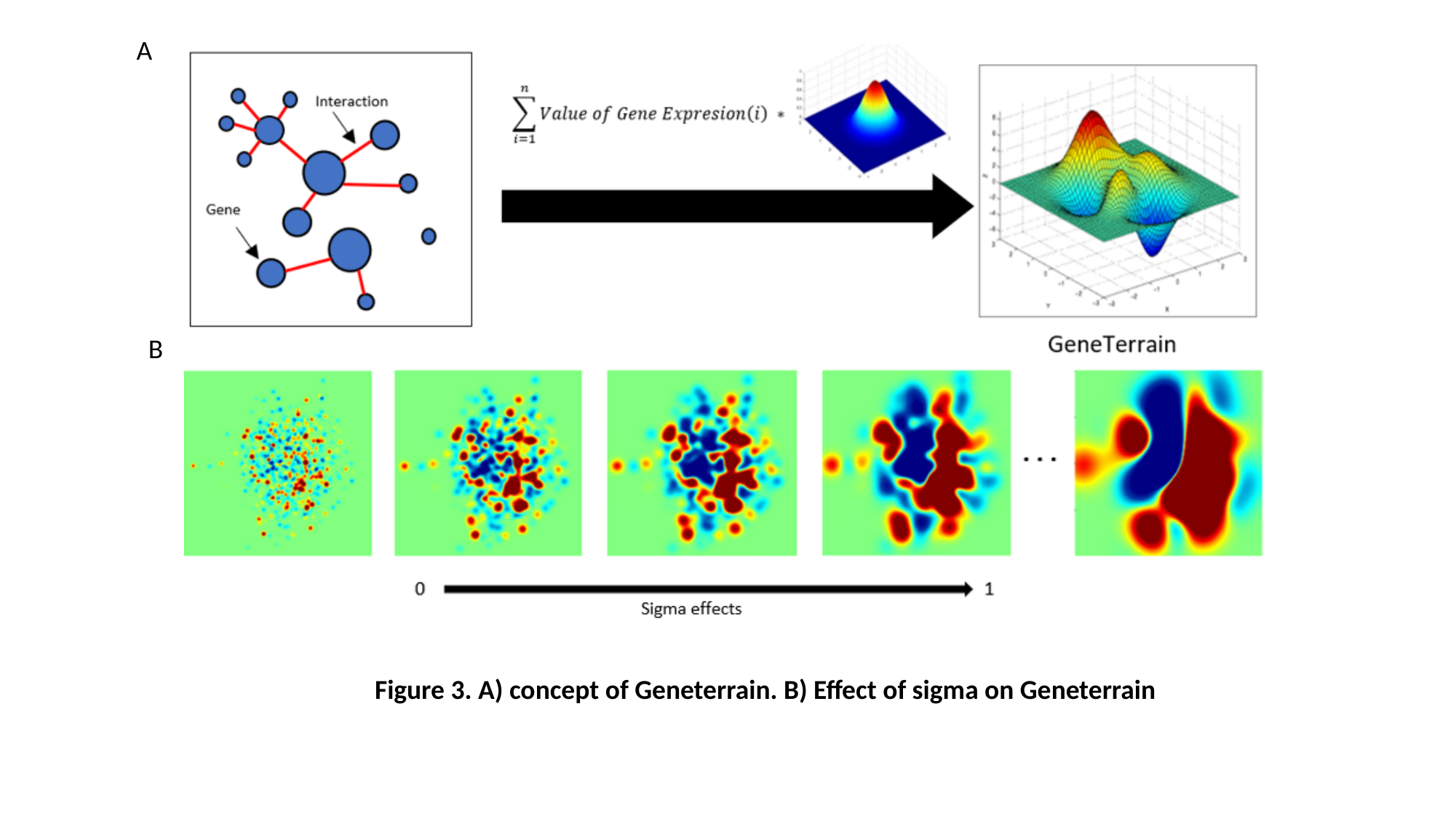

A
B
Figure 3. A) concept of Geneterrain. B) Effect of sigma on Geneterrain
