## supplementary for "Explorative Discovery of Gene Signatures and Clinotypes in Glioblastoma Cancer Through GeneTerrain Knowledge Map Representation": Supplementary 2.pptx

### Slide 1
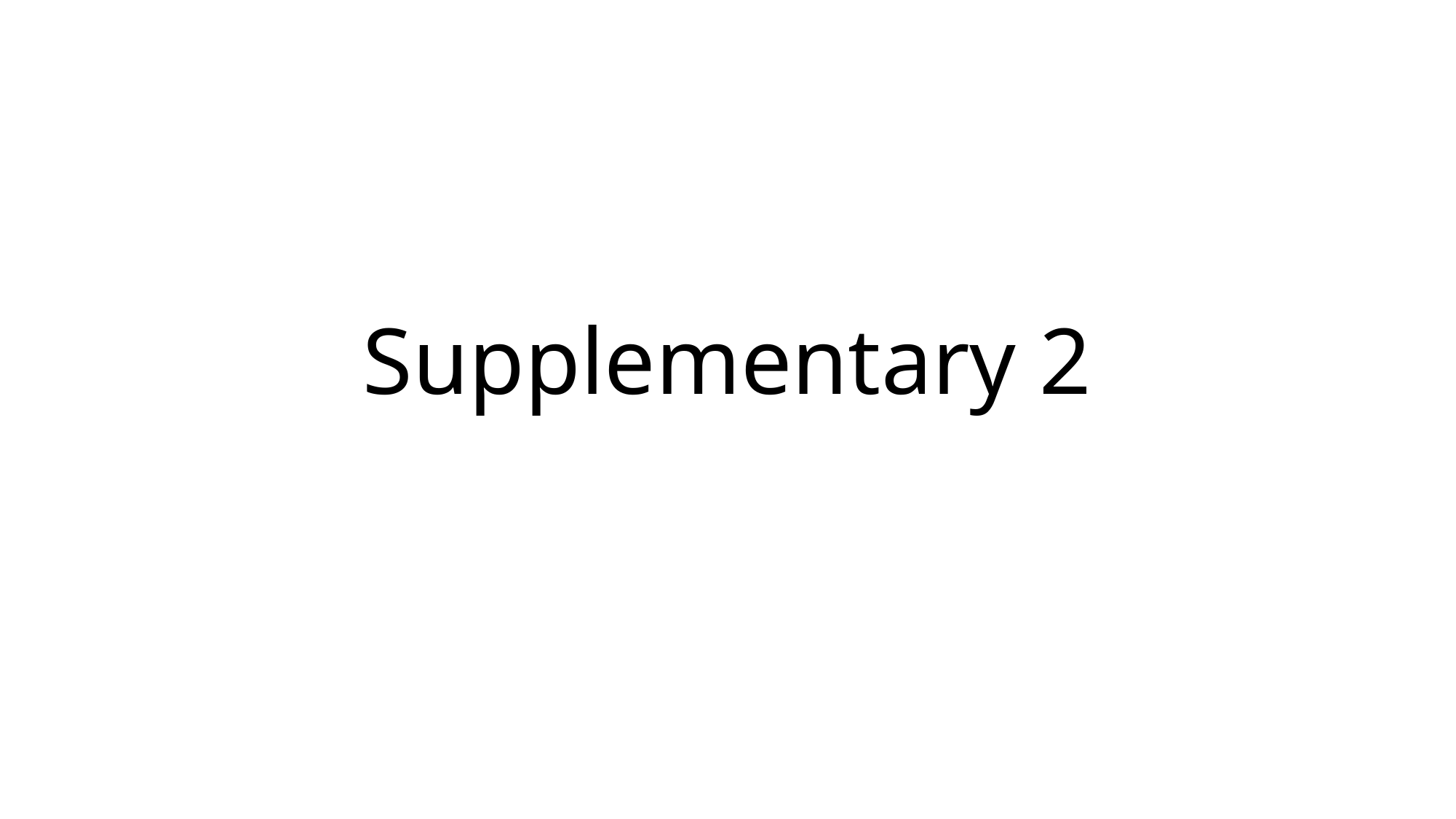

#### Supplementary 2

### Slide 2
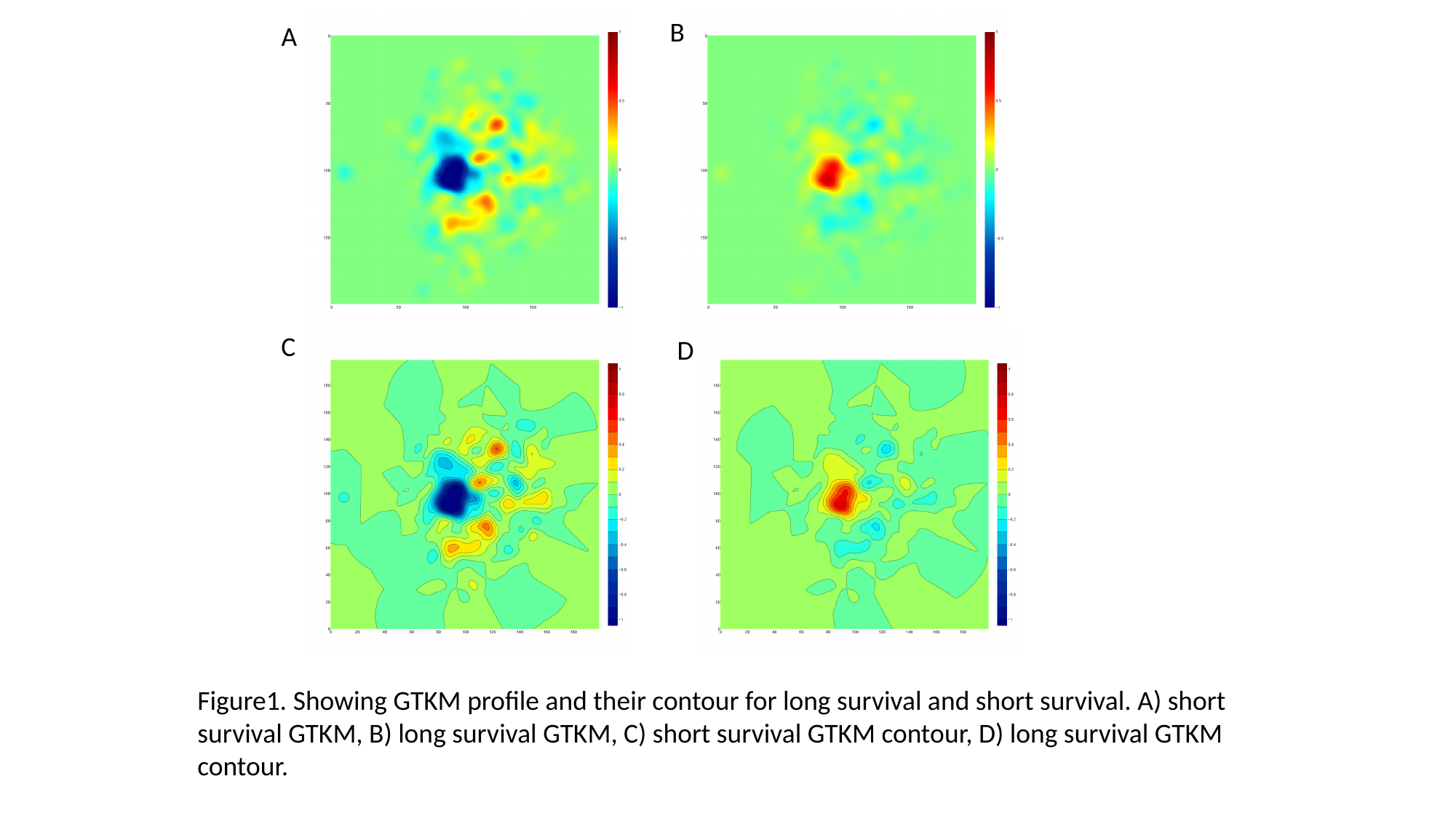

B
A
C
D
Figure1. Showing GTKM profile and their contour for long survival and short survival. A) short survival GTKM, B) long survival GTKM, C) short survival GTKM contour, D) long survival GTKM contour.

### Slide 3
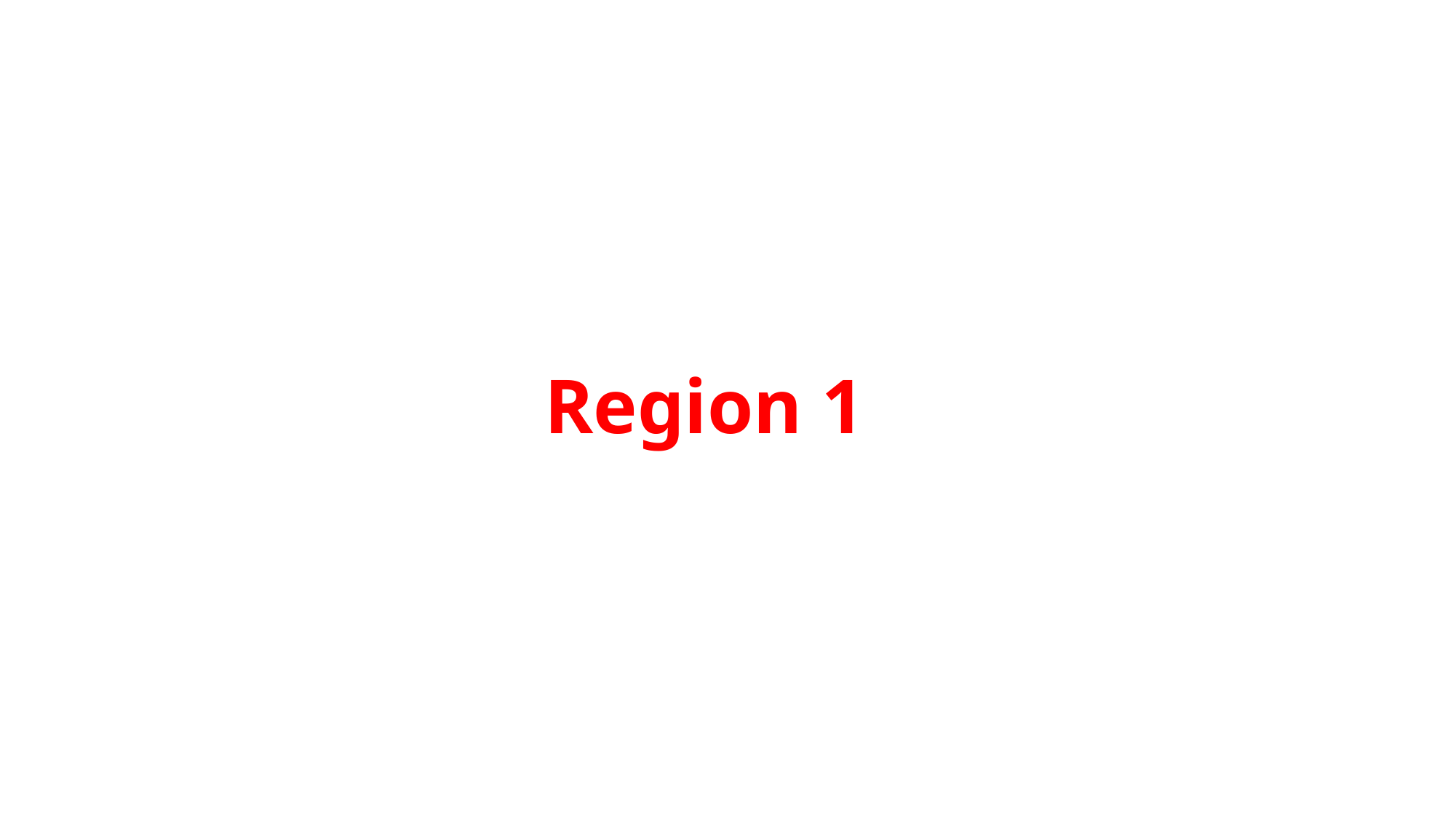

#### Region 1

### Slide 4
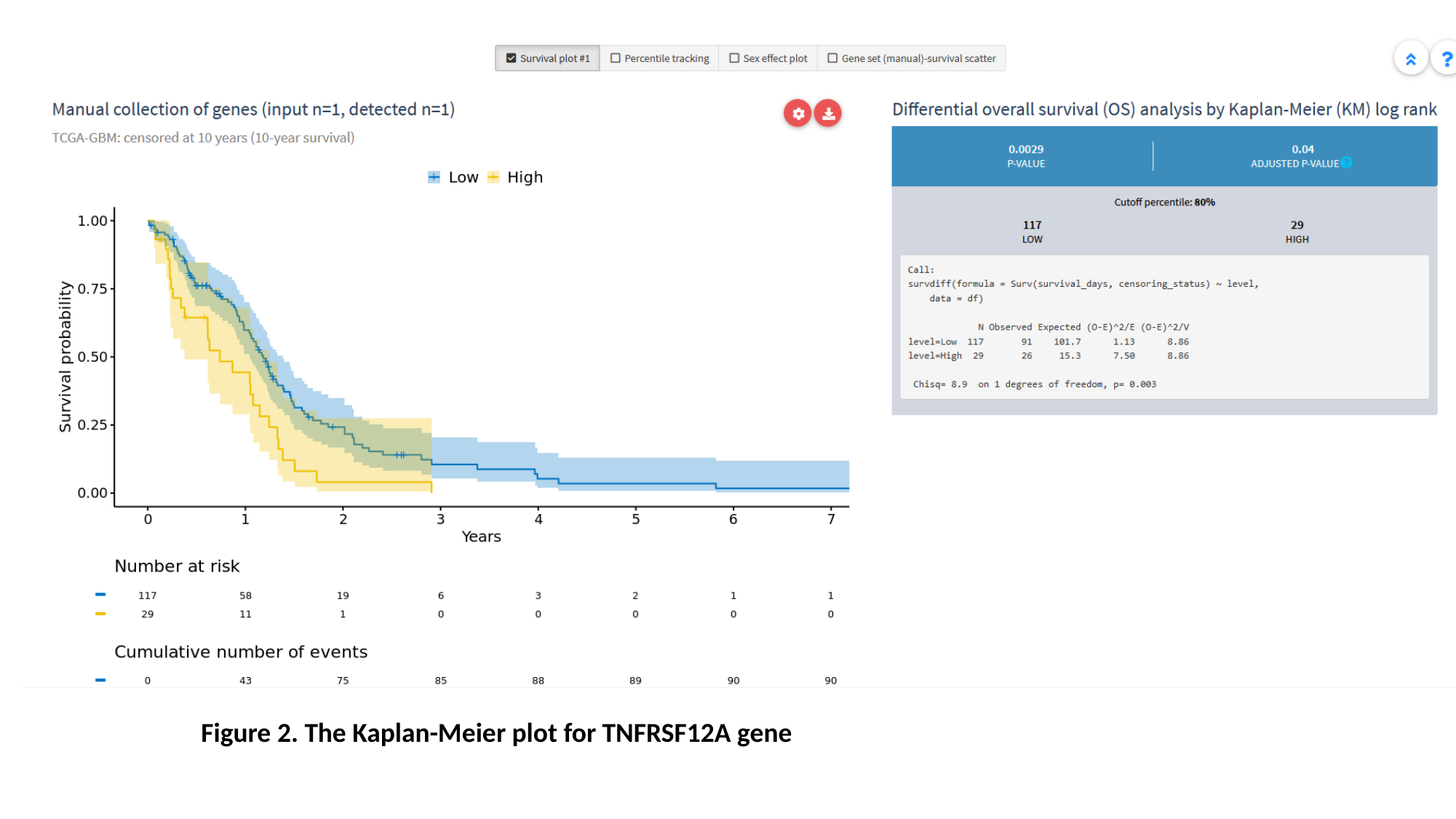

Figure 2. The Kaplan-Meier plot for TNFRSF12A gene

### Slide 5
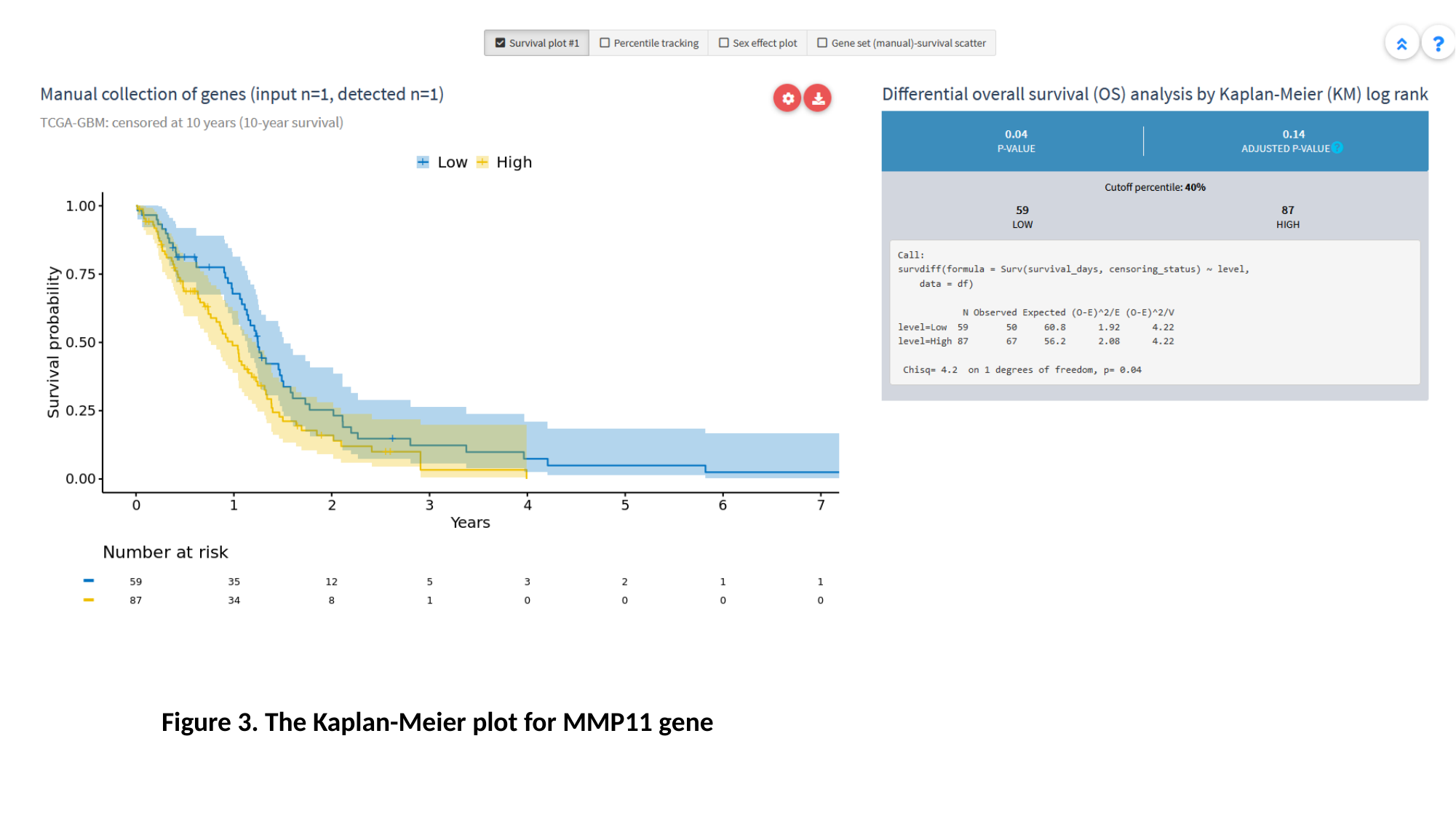

Figure 3. The Kaplan-Meier plot for MMP11 gene

### Slide 6
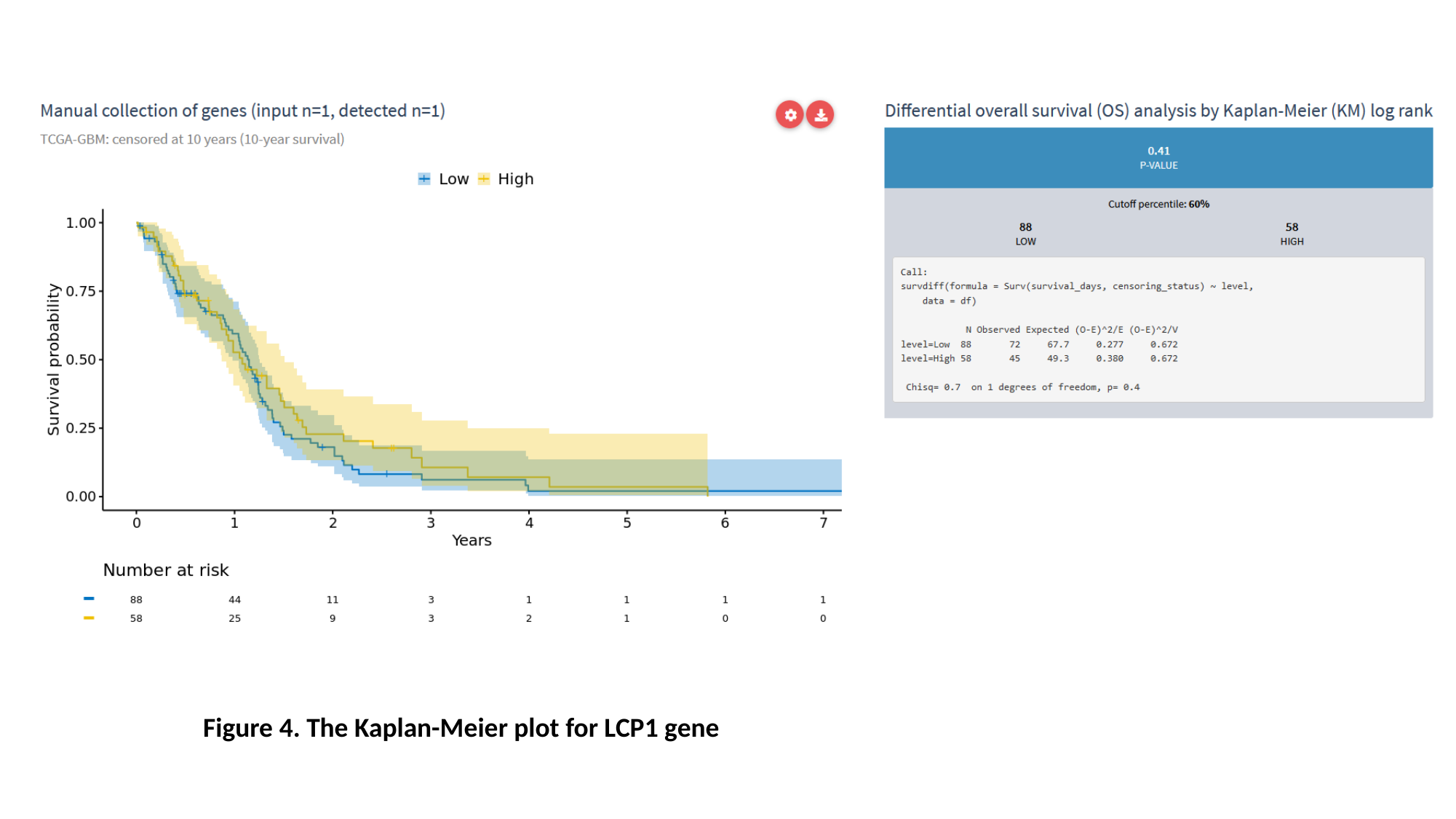

#### Figure 4. The Kaplan-Meier plot for LCP1 gene

### Slide 7
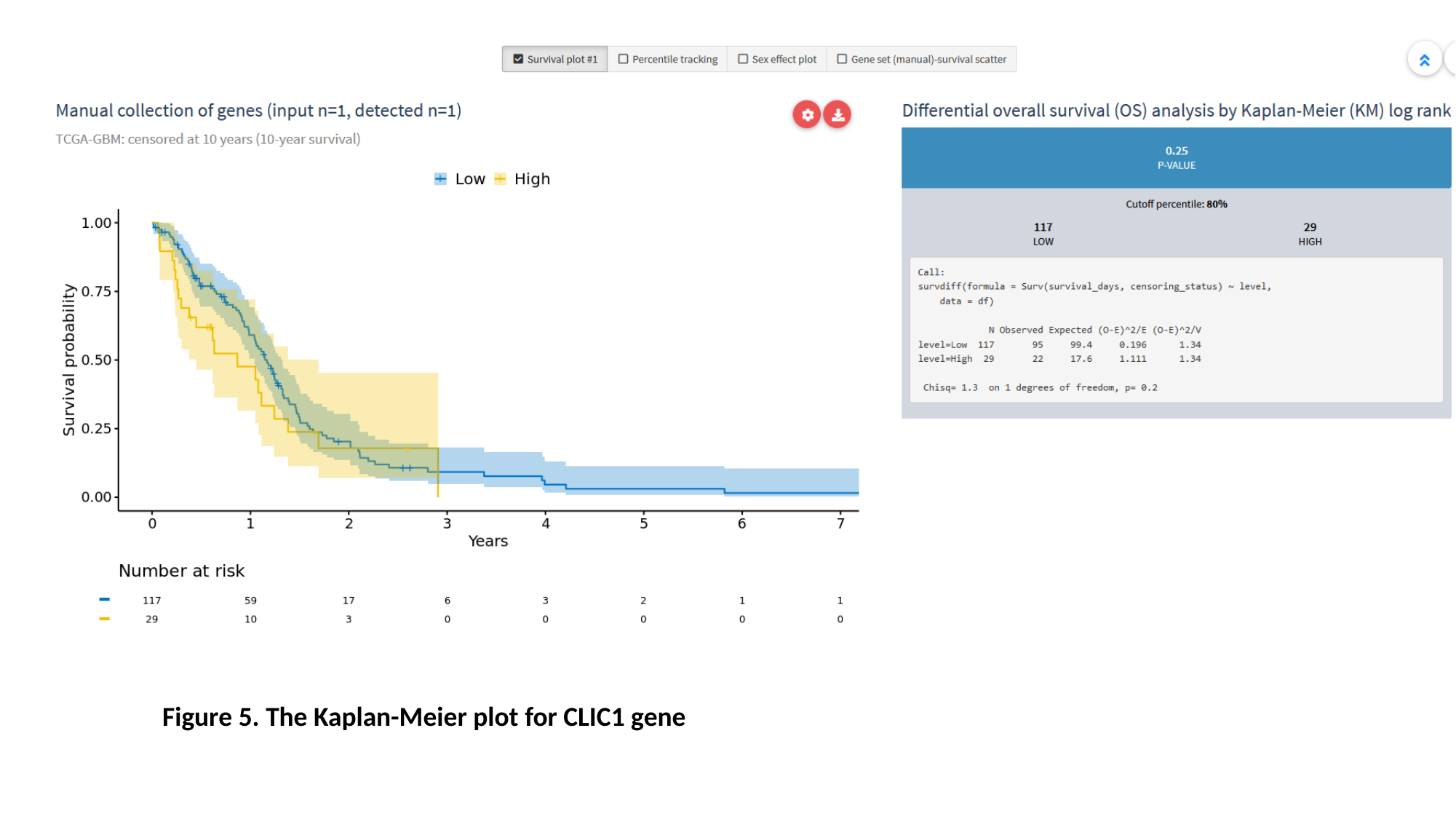

#### Figure 5. The Kaplan-Meier plot for CLIC1 gene

### Slide 8
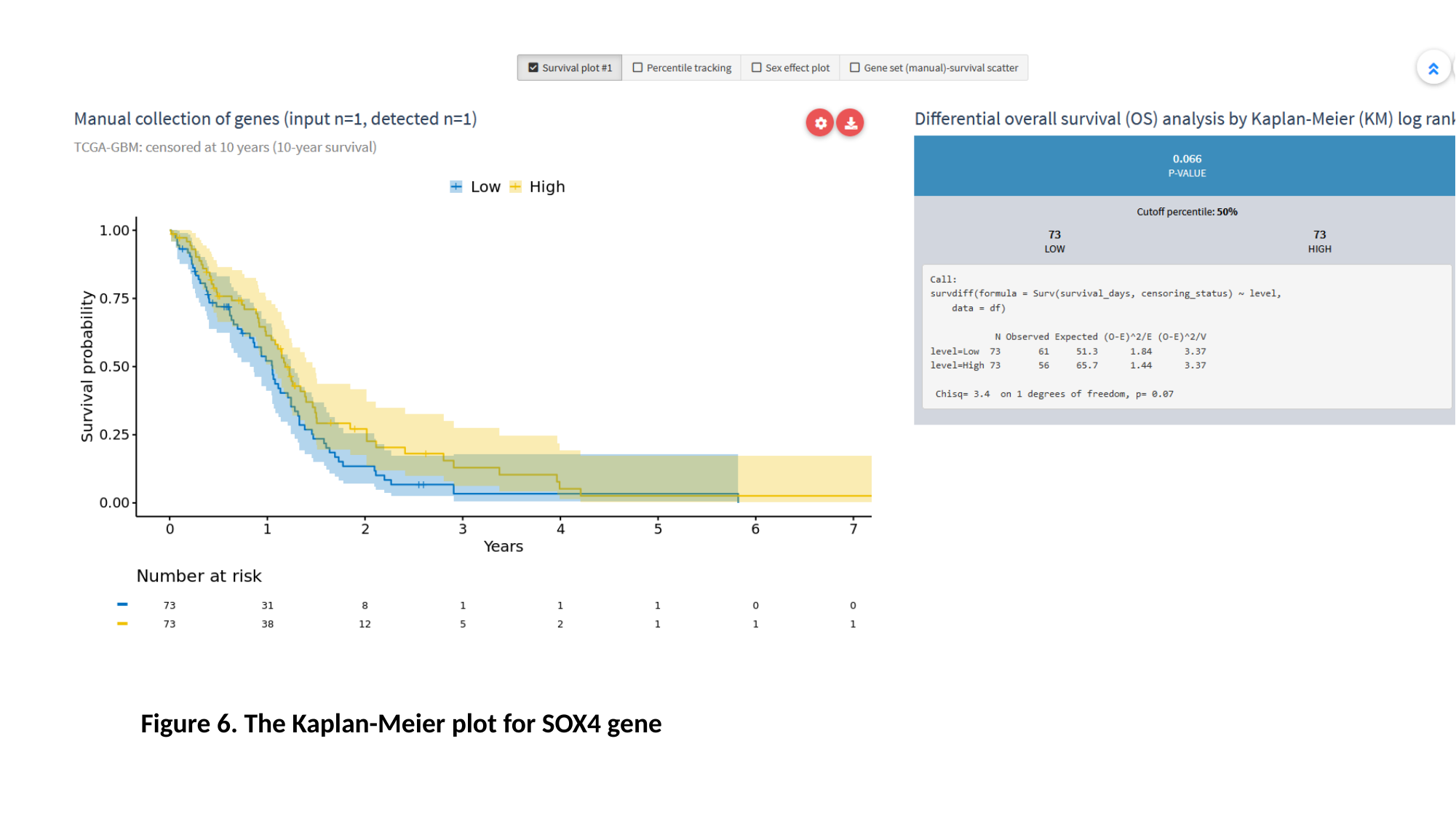

#### Figure 6. The Kaplan-Meier plot for SOX4 gene

### Slide 9
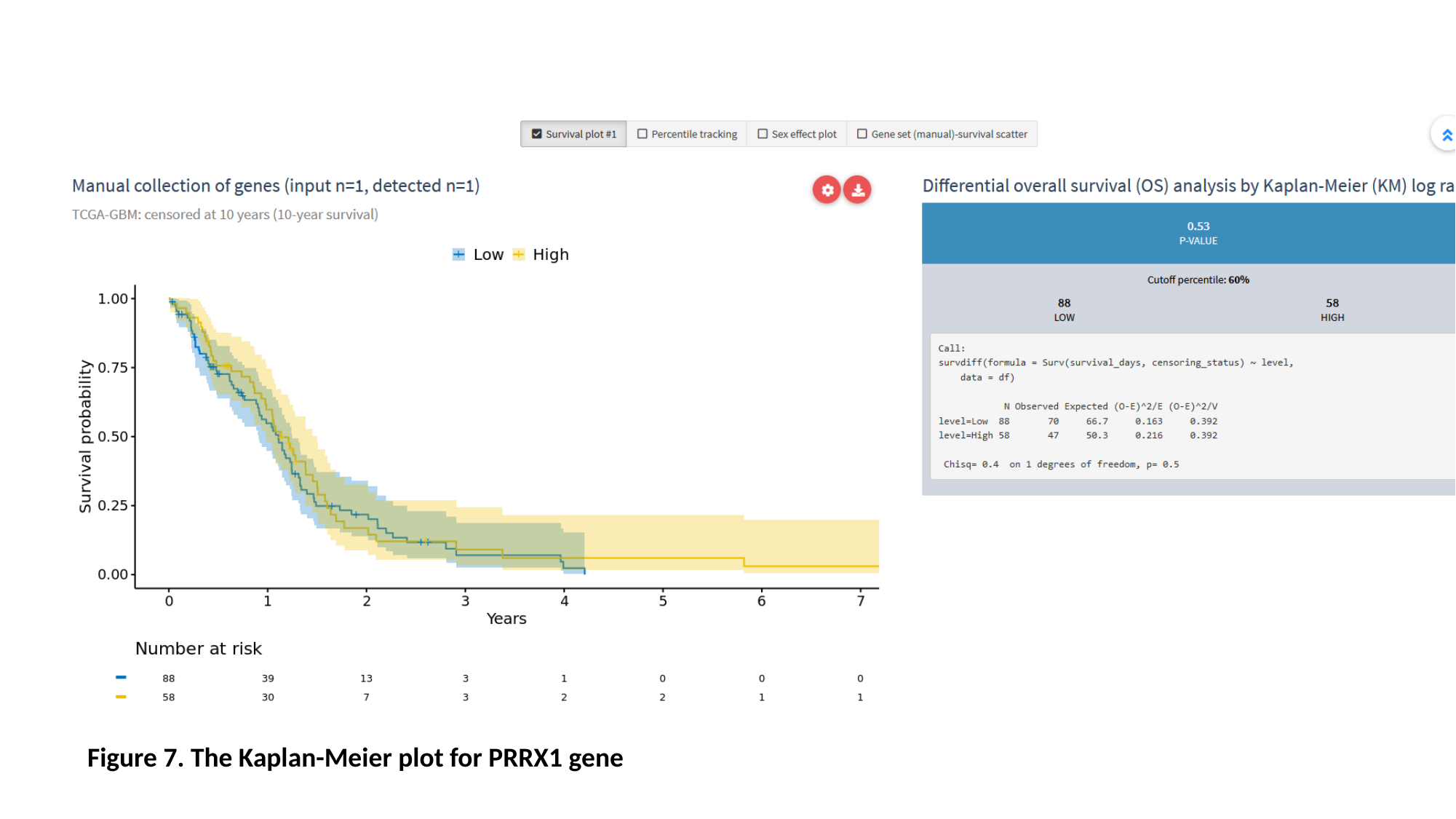

#### Figure 7. The Kaplan-Meier plot for PRRX1 gene

### Slide 10
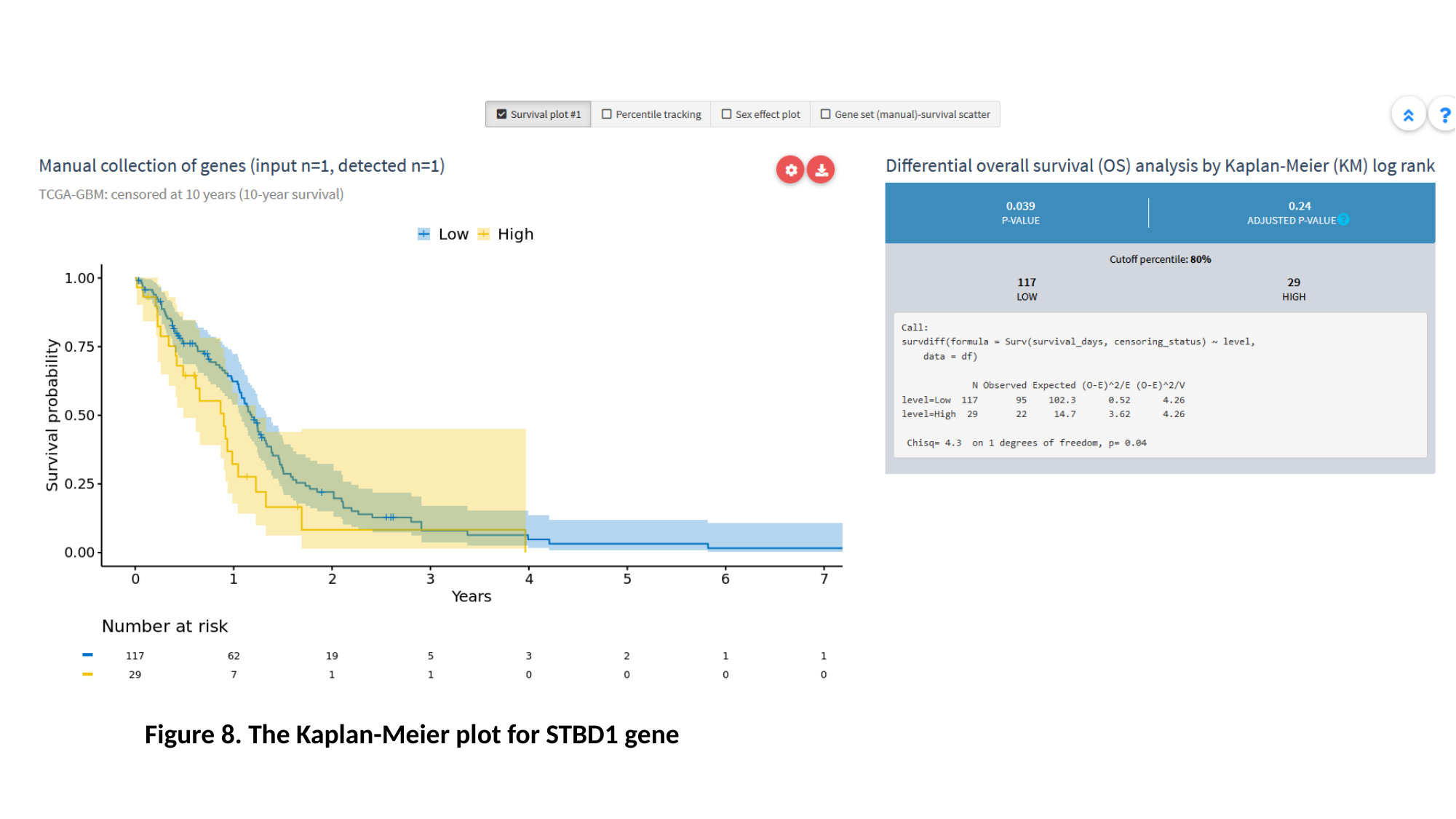

#### Figure 8. The Kaplan-Meier plot for STBD1 gene

### Slide 11
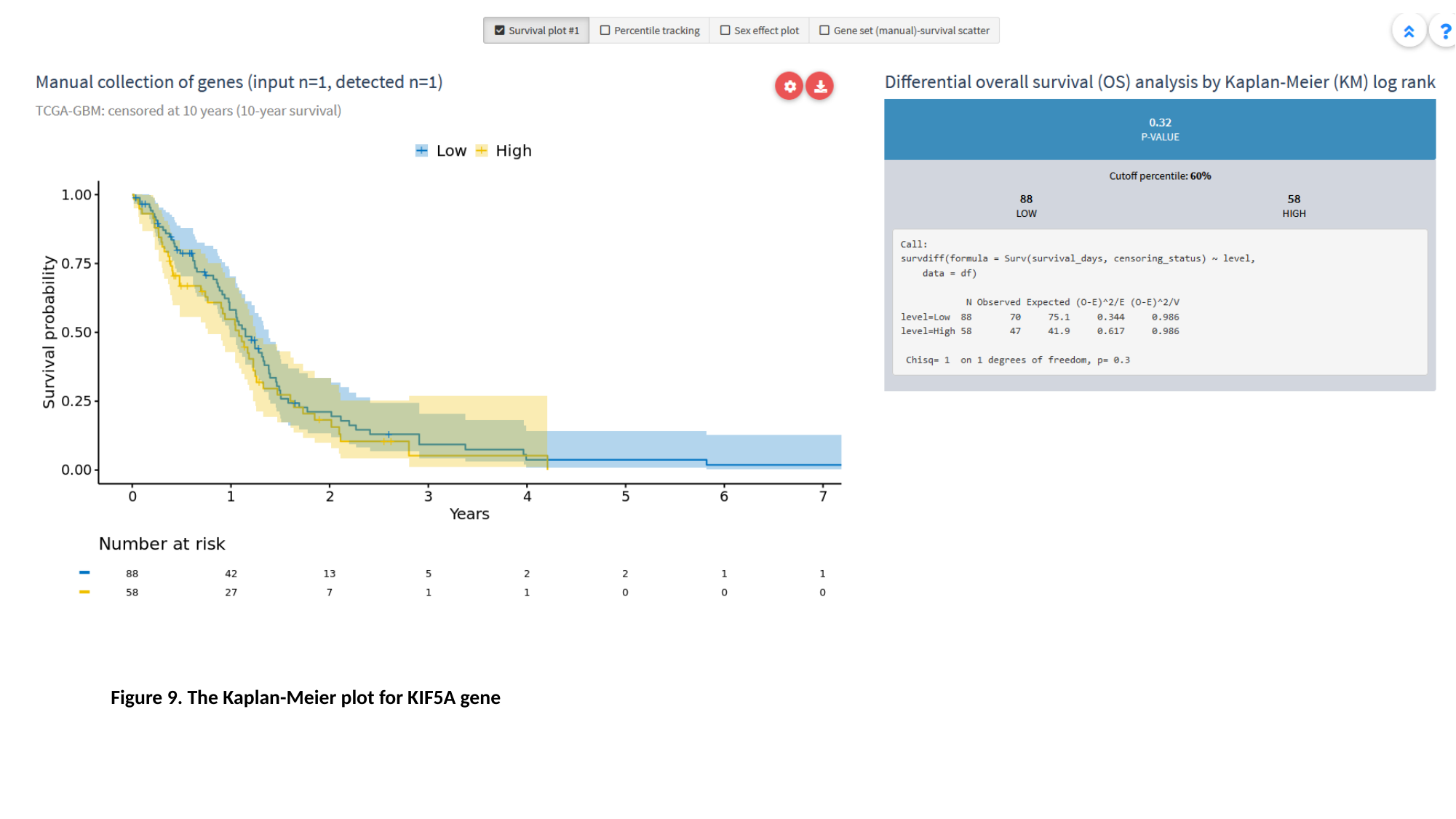

#### Figure 9. The Kaplan-Meier plot for KIF5A gene

### Slide 12
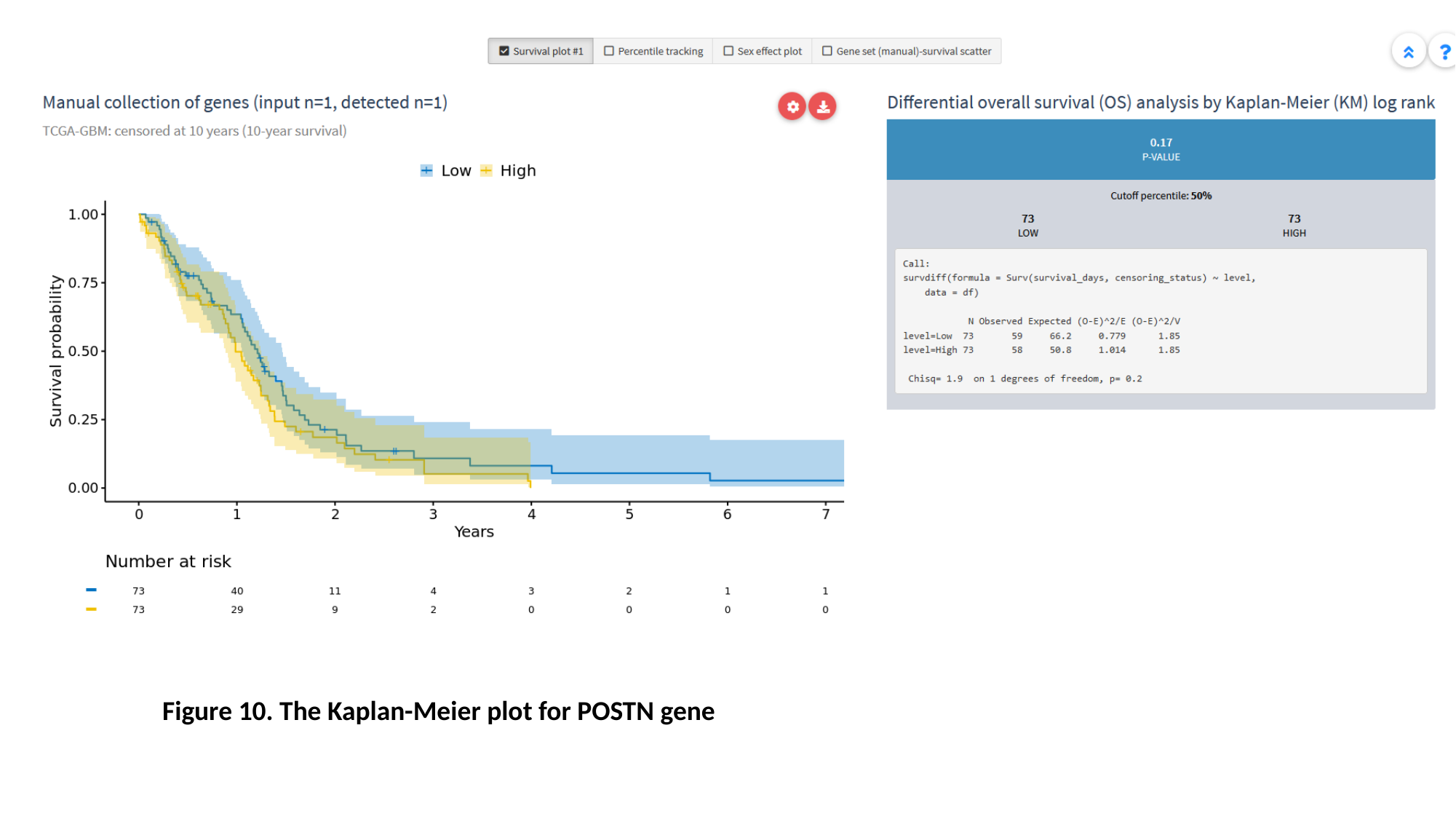

#### Figure 10. The Kaplan-Meier plot for POSTN gene

### Slide 13
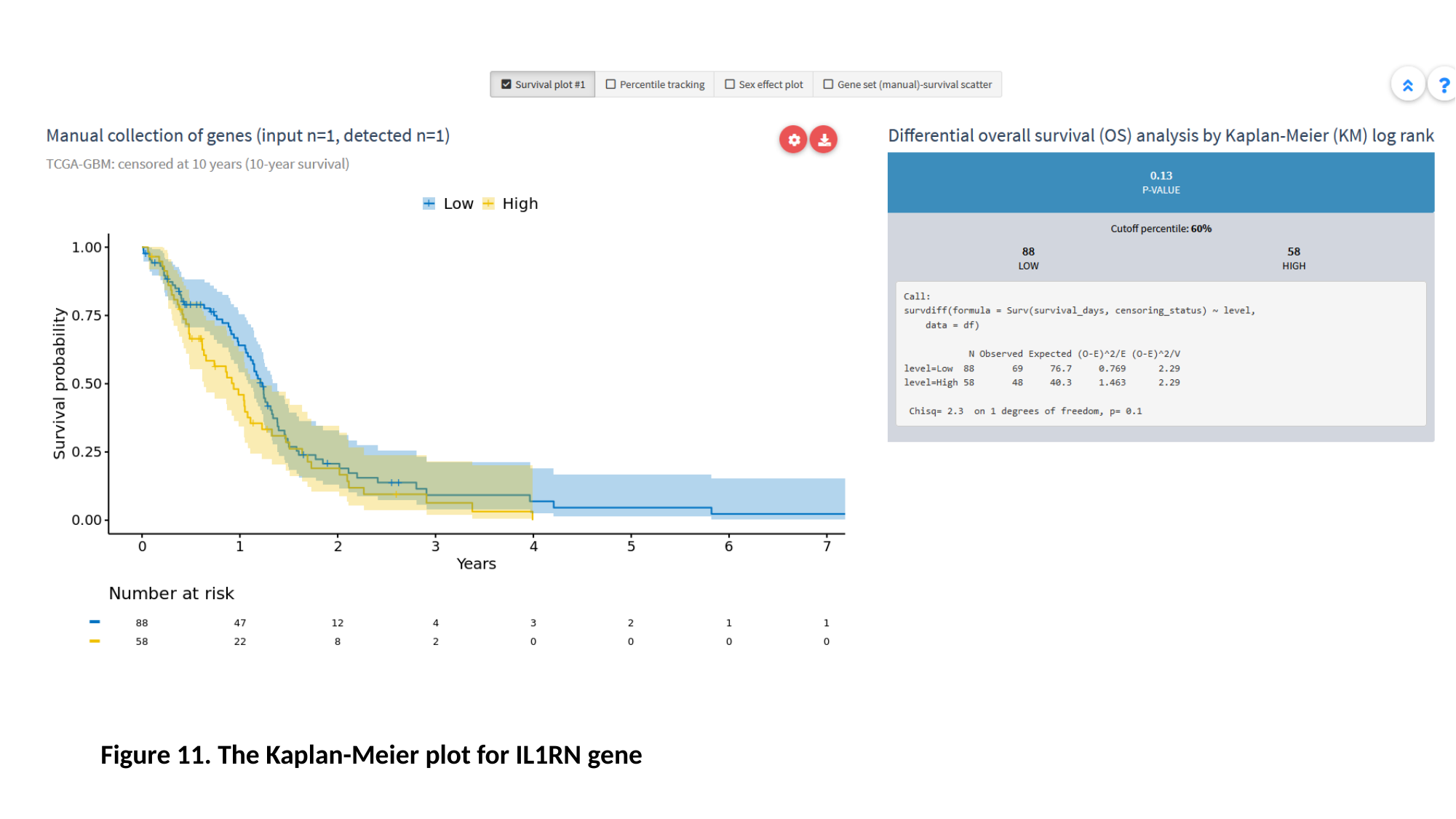

#### Figure 11. The Kaplan-Meier plot for IL1RN gene

### Slide 14
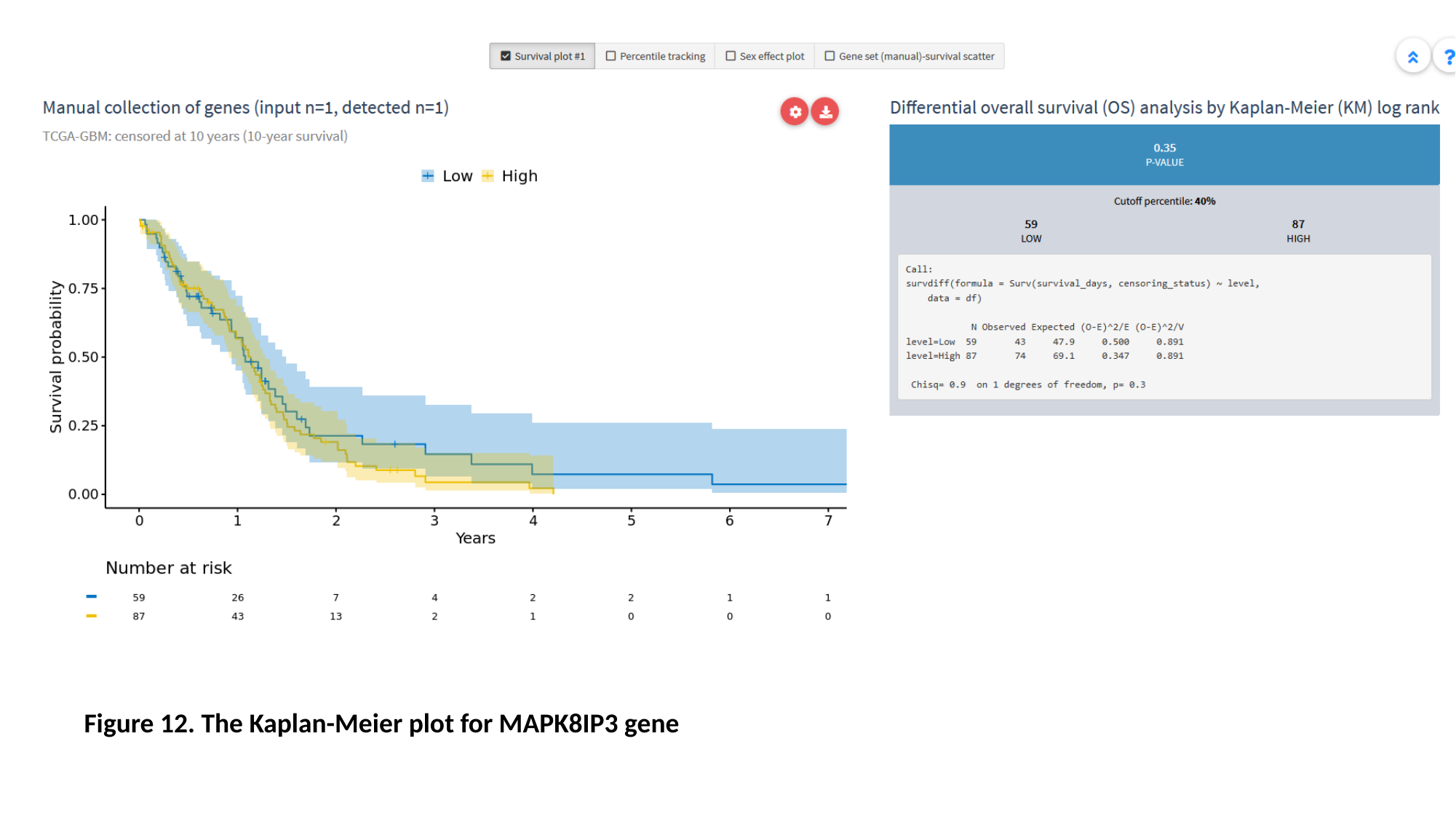

#### Figure 12. The Kaplan-Meier plot for MAPK8IP3 gene

### Slide 15
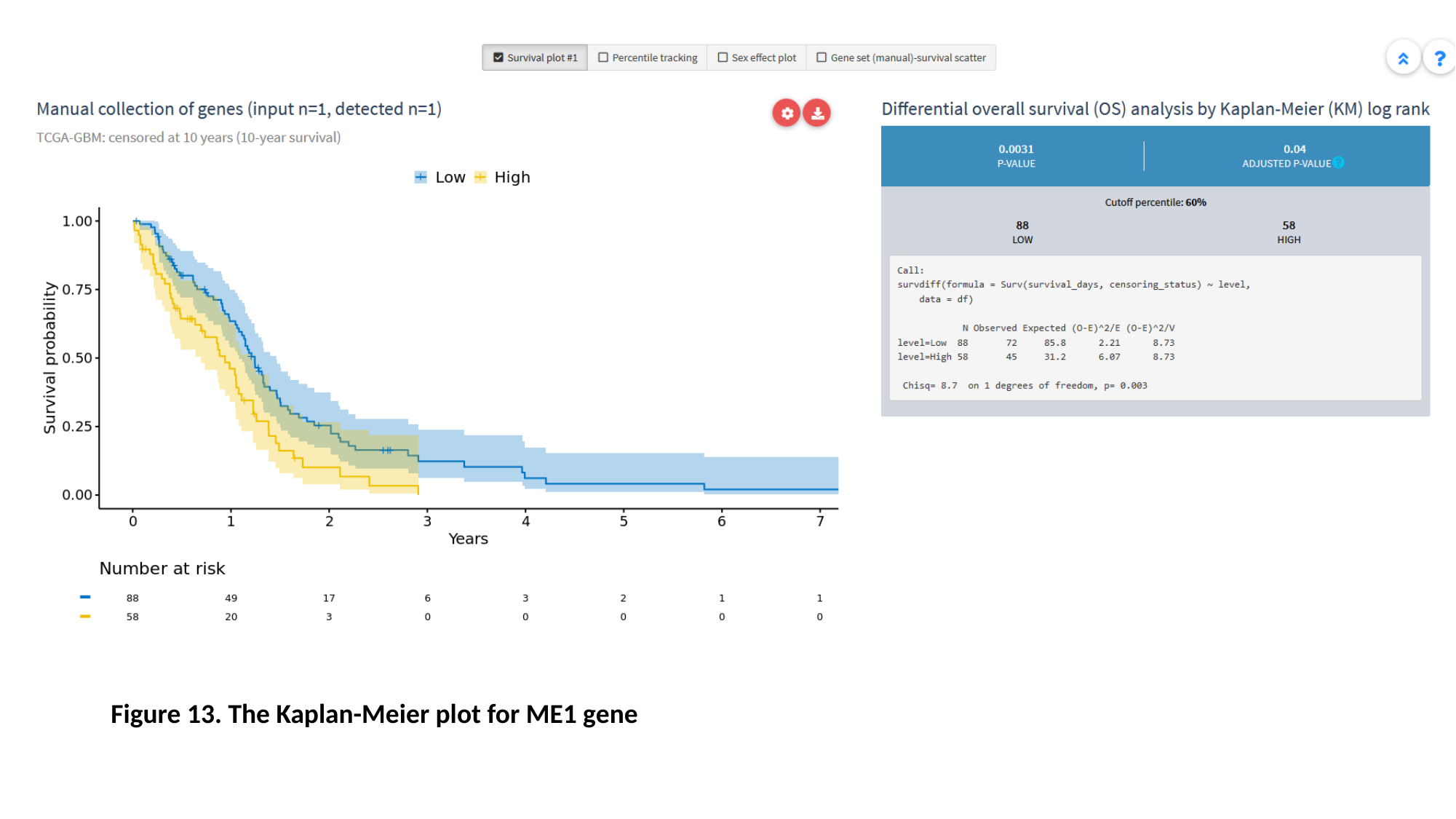

Figure 13. The Kaplan-Meier plot for ME1 gene

### Slide 16
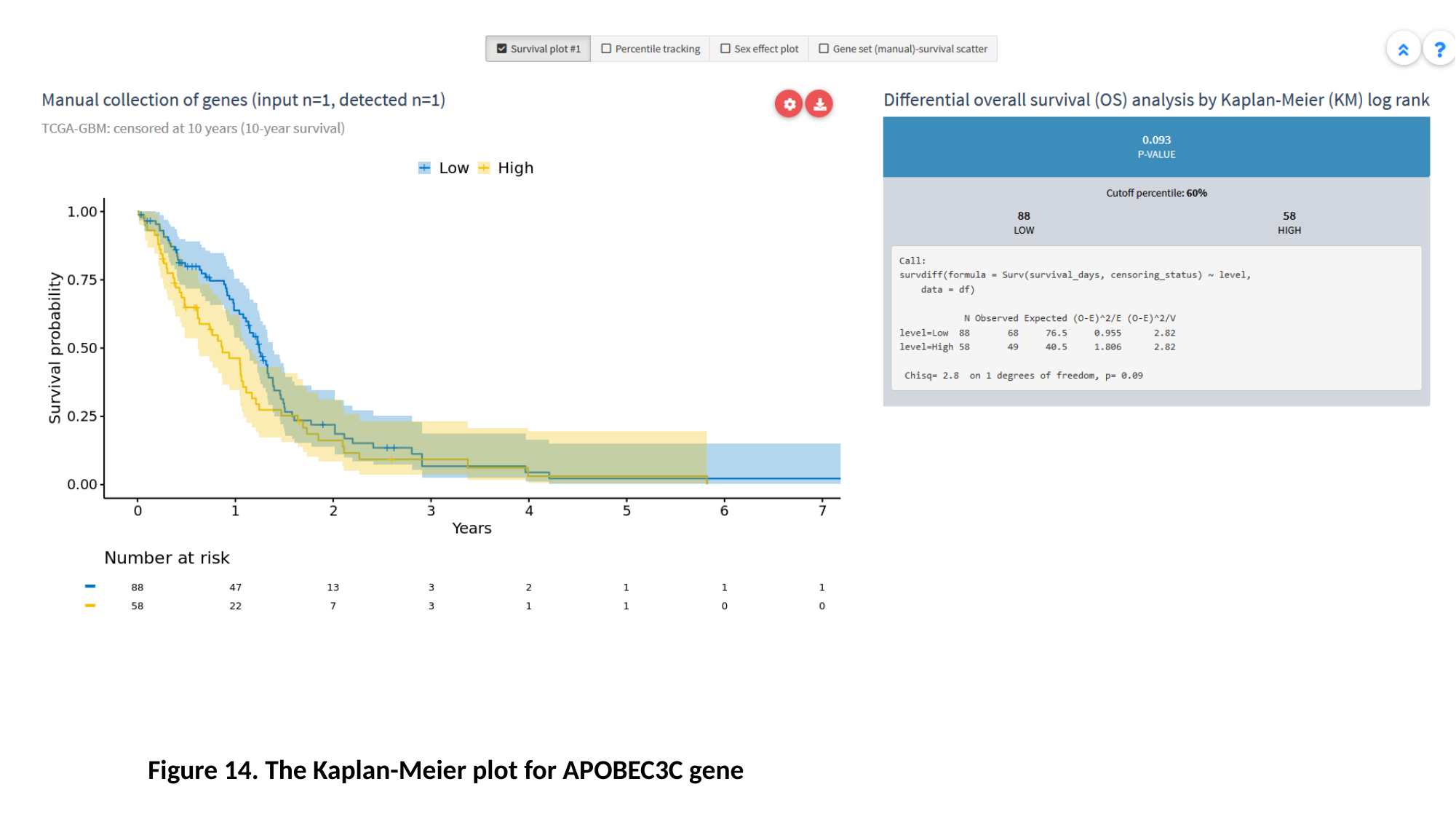

Figure 14. The Kaplan-Meier plot for APOBEC3C gene

### Slide 17
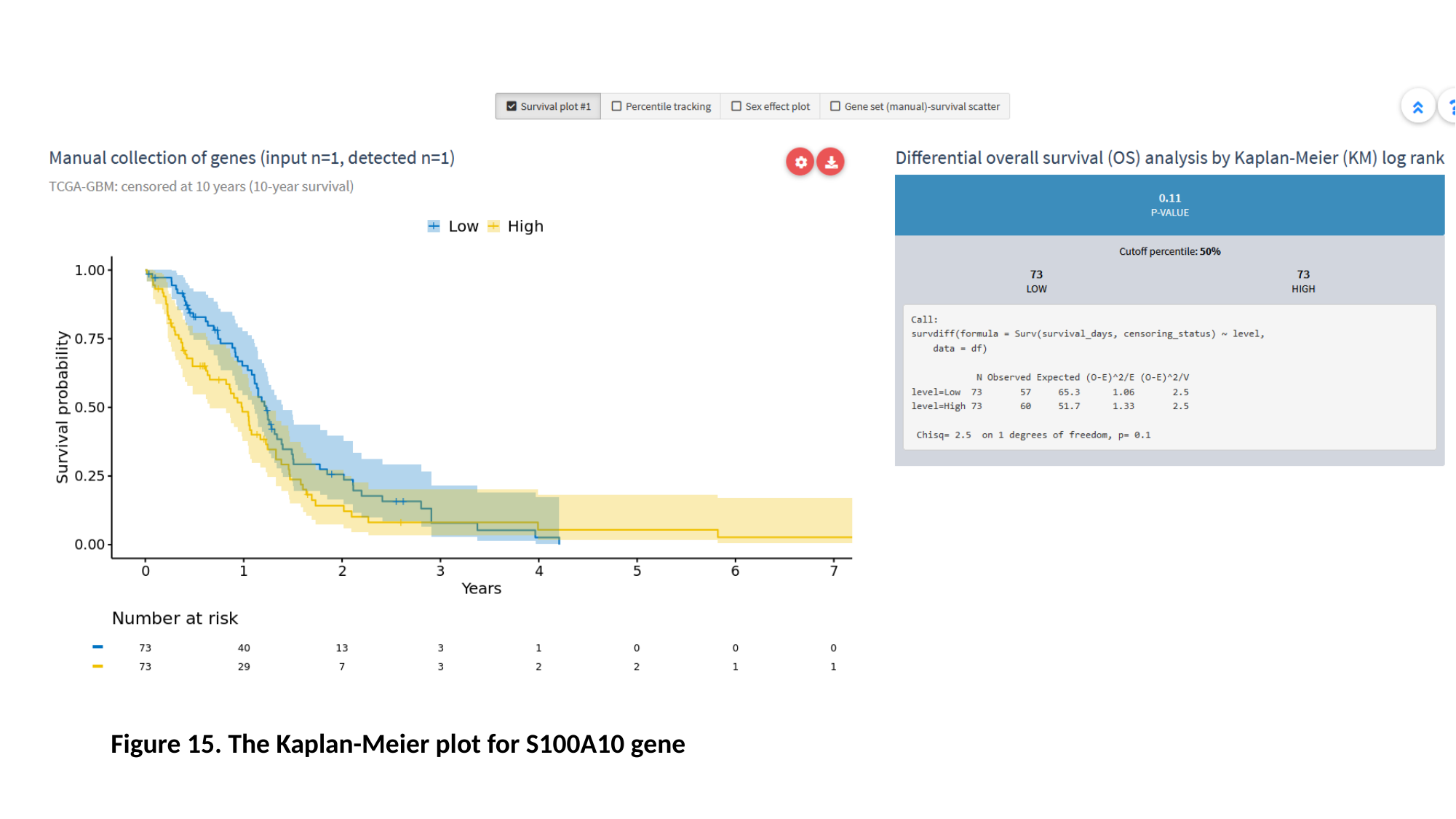

Figure 15. The Kaplan-Meier plot for S100A10 gene

### Slide 18
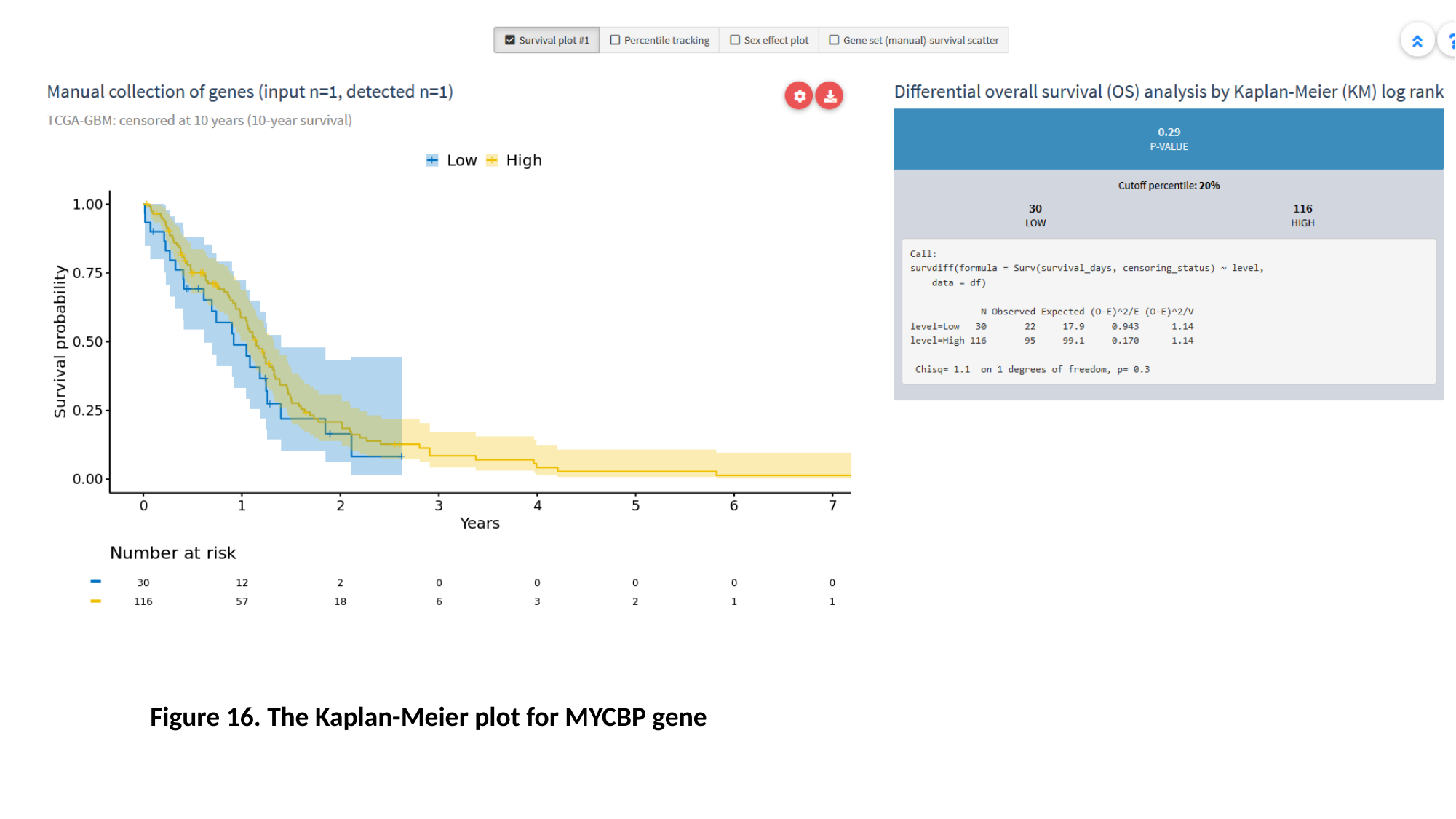

Figure 16. The Kaplan-Meier plot for MYCBP gene

### Slide 19
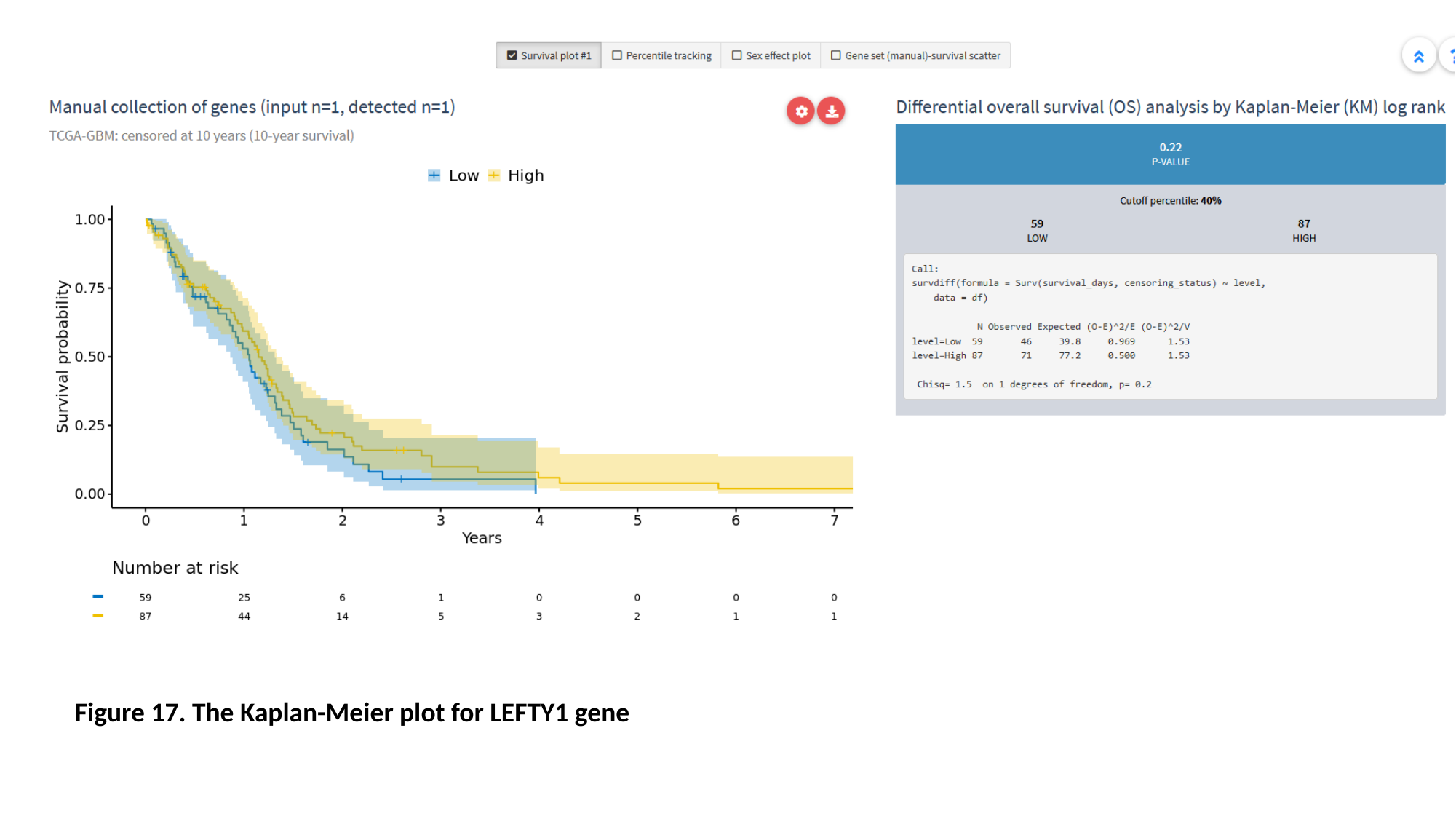

Figure 17. The Kaplan-Meier plot for LEFTY1 gene

### Slide 20
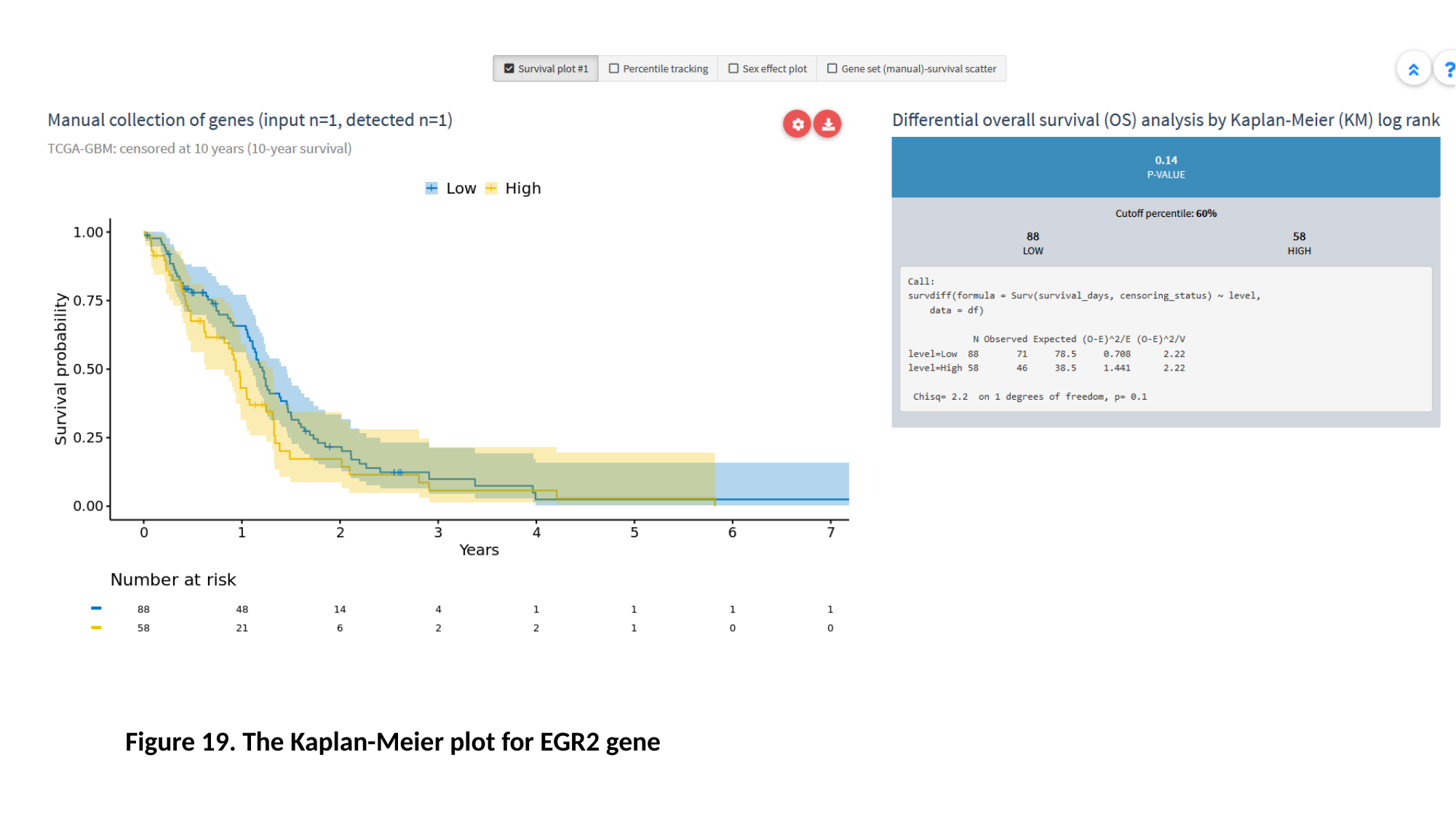

Figure 19. The Kaplan-Meier plot for EGR2 gene

### Slide 21
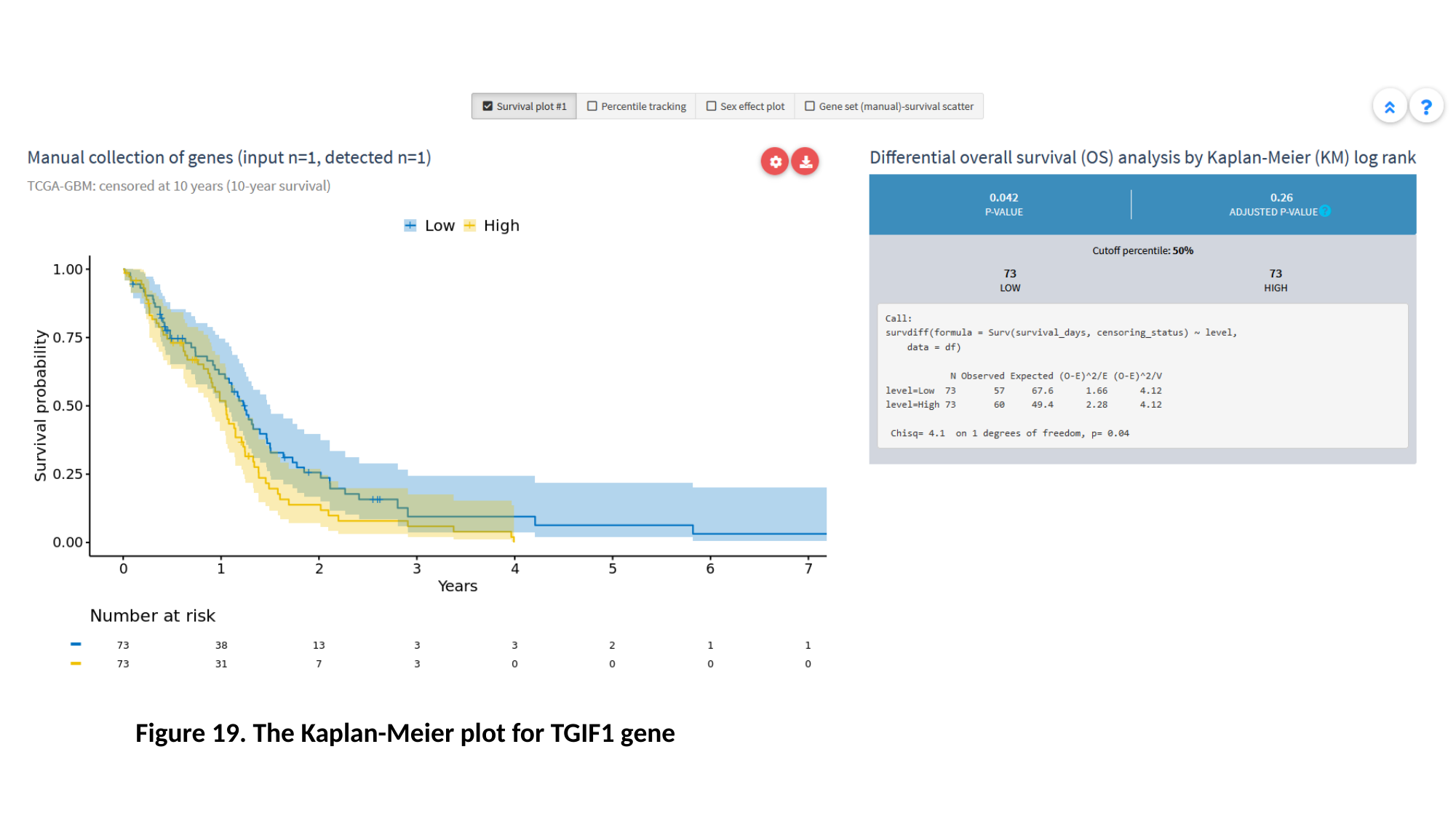

Figure 19. The Kaplan-Meier plot for TGIF1 gene

### Slide 22
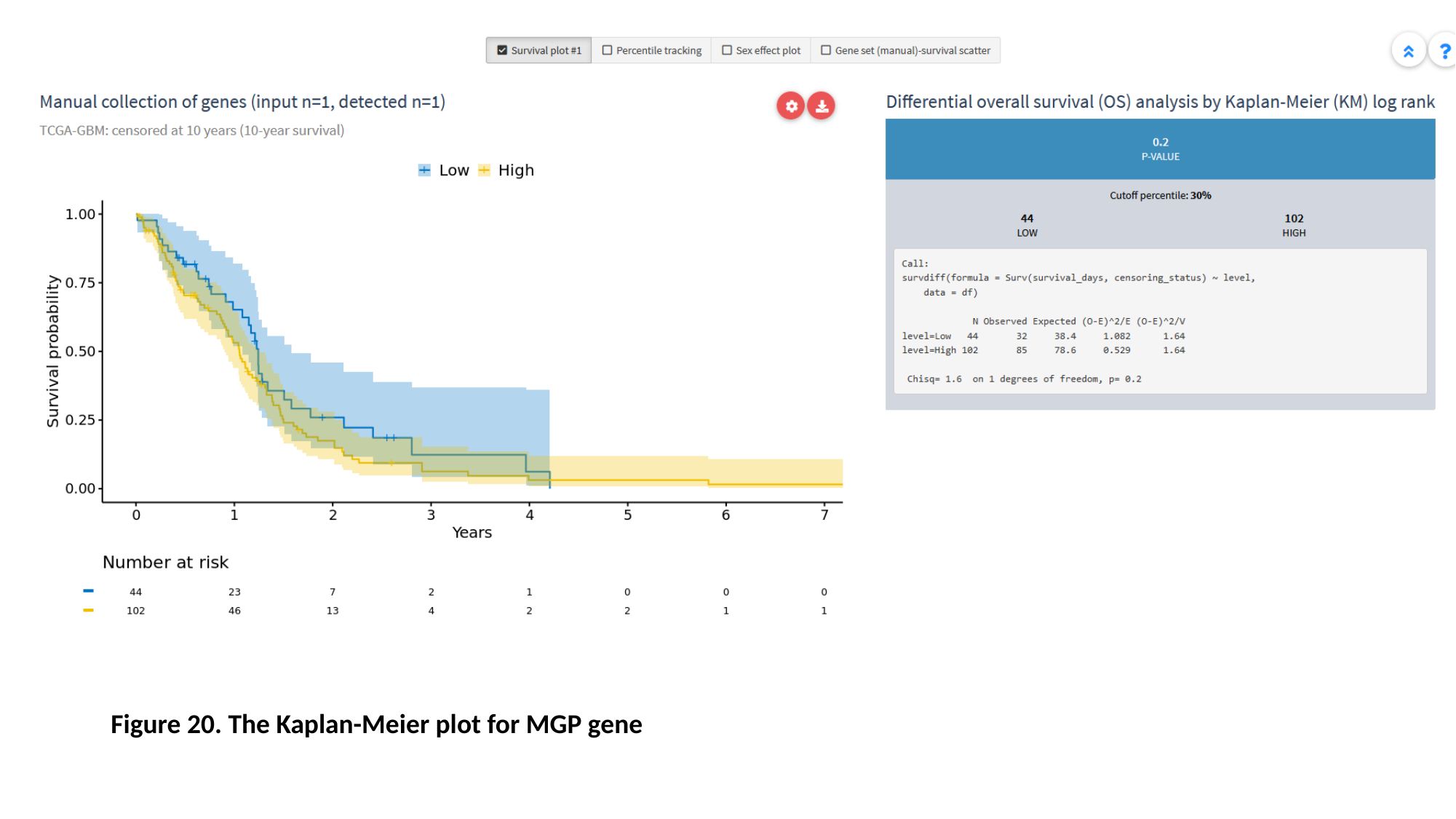

Figure 20. The Kaplan-Meier plot for MGP gene

### Slide 23
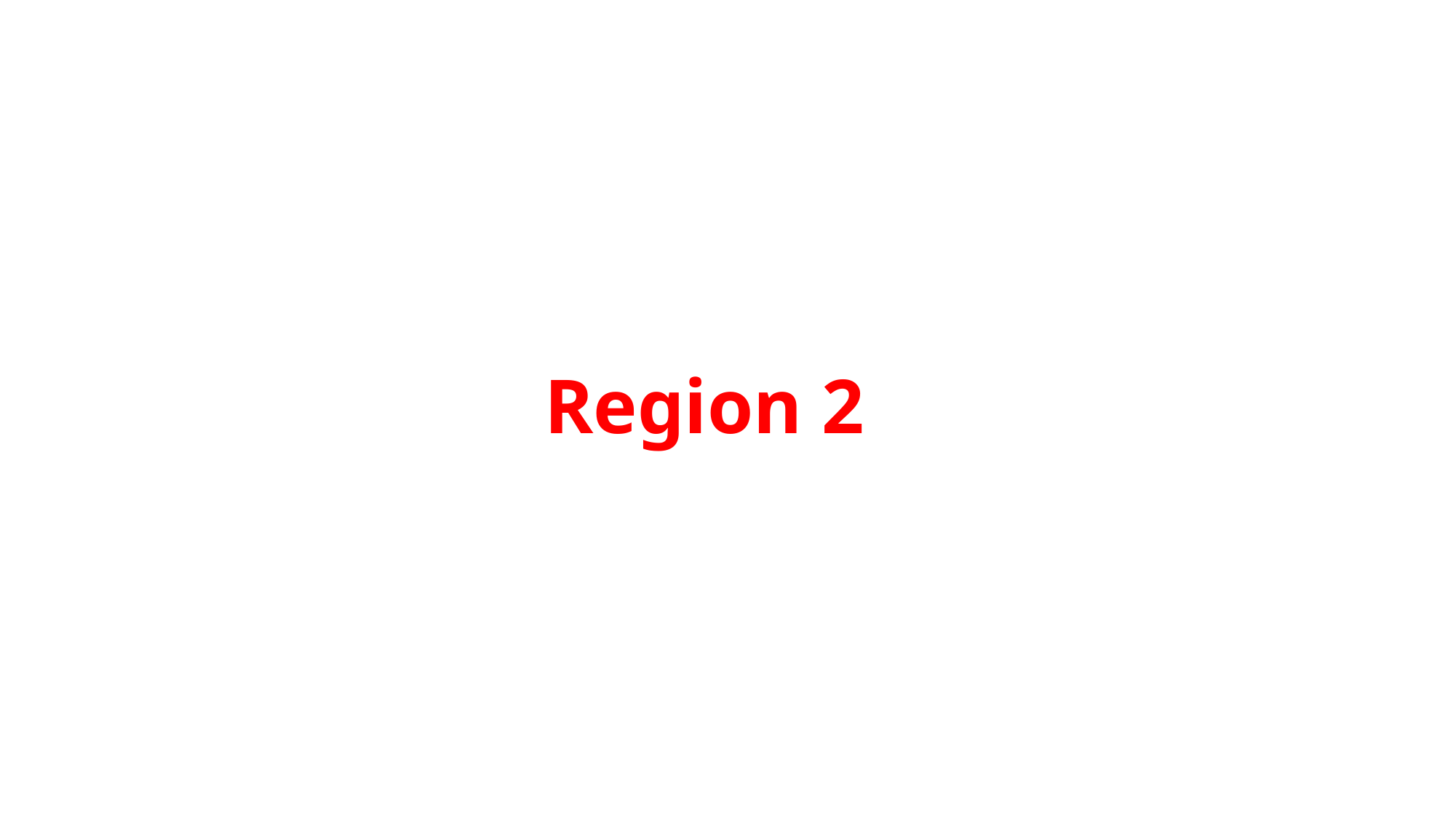

#### Region 2

### Slide 24
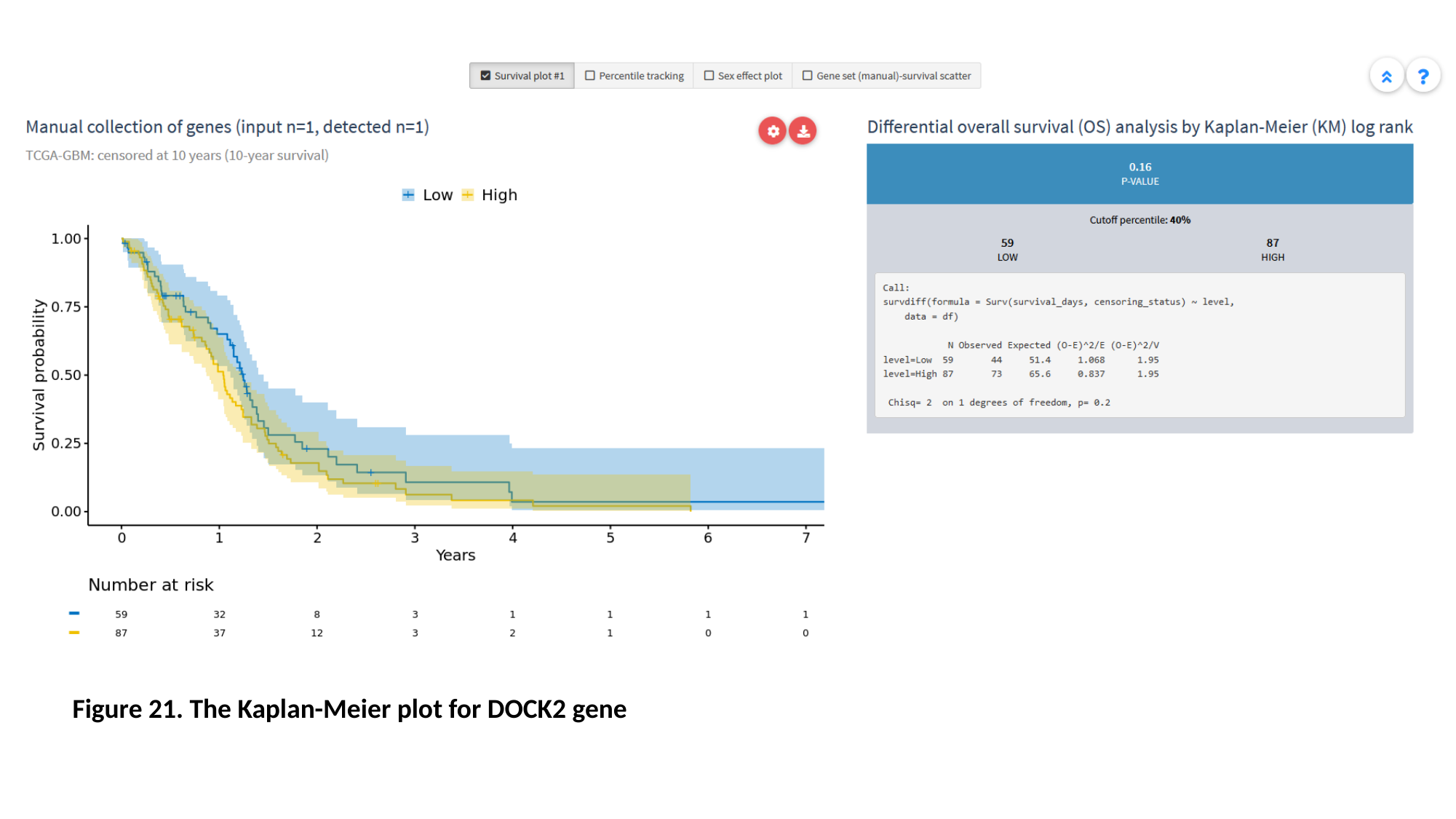

Figure 21. The Kaplan-Meier plot for DOCK2 gene

### Slide 25
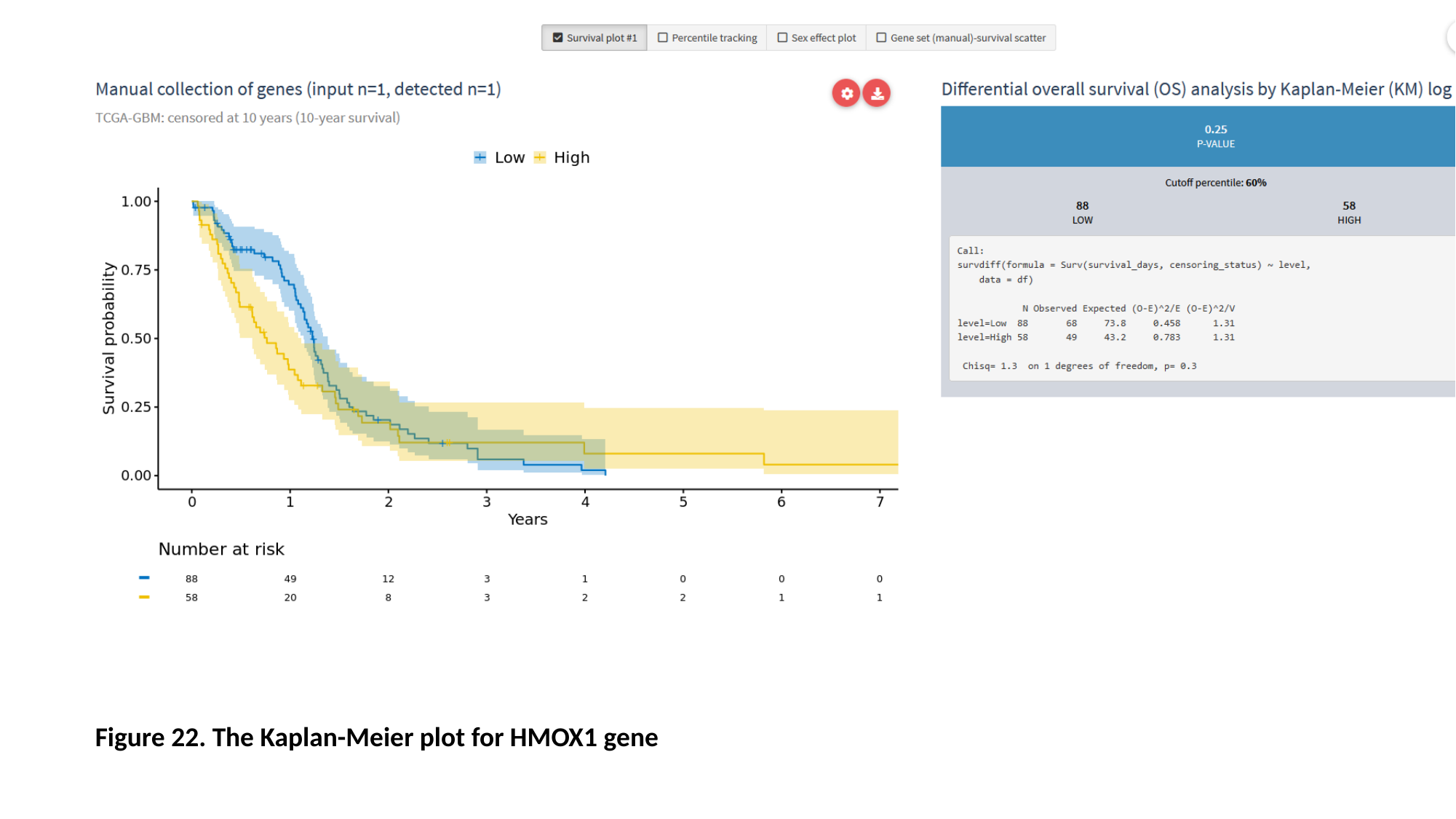

Figure 22. The Kaplan-Meier plot for HMOX1 gene

### Slide 26
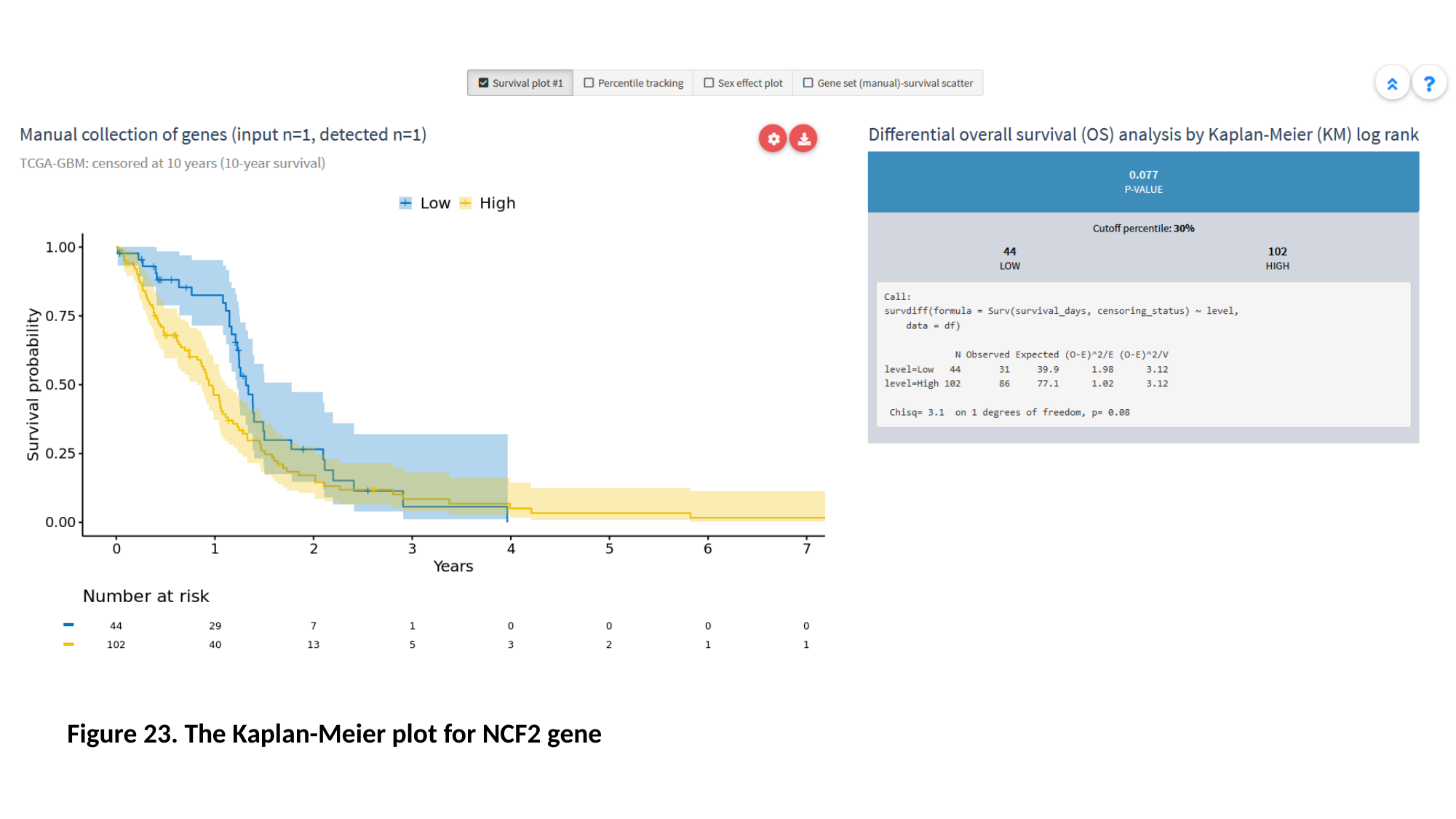

Figure 23. The Kaplan-Meier plot for NCF2 gene

### Slide 27
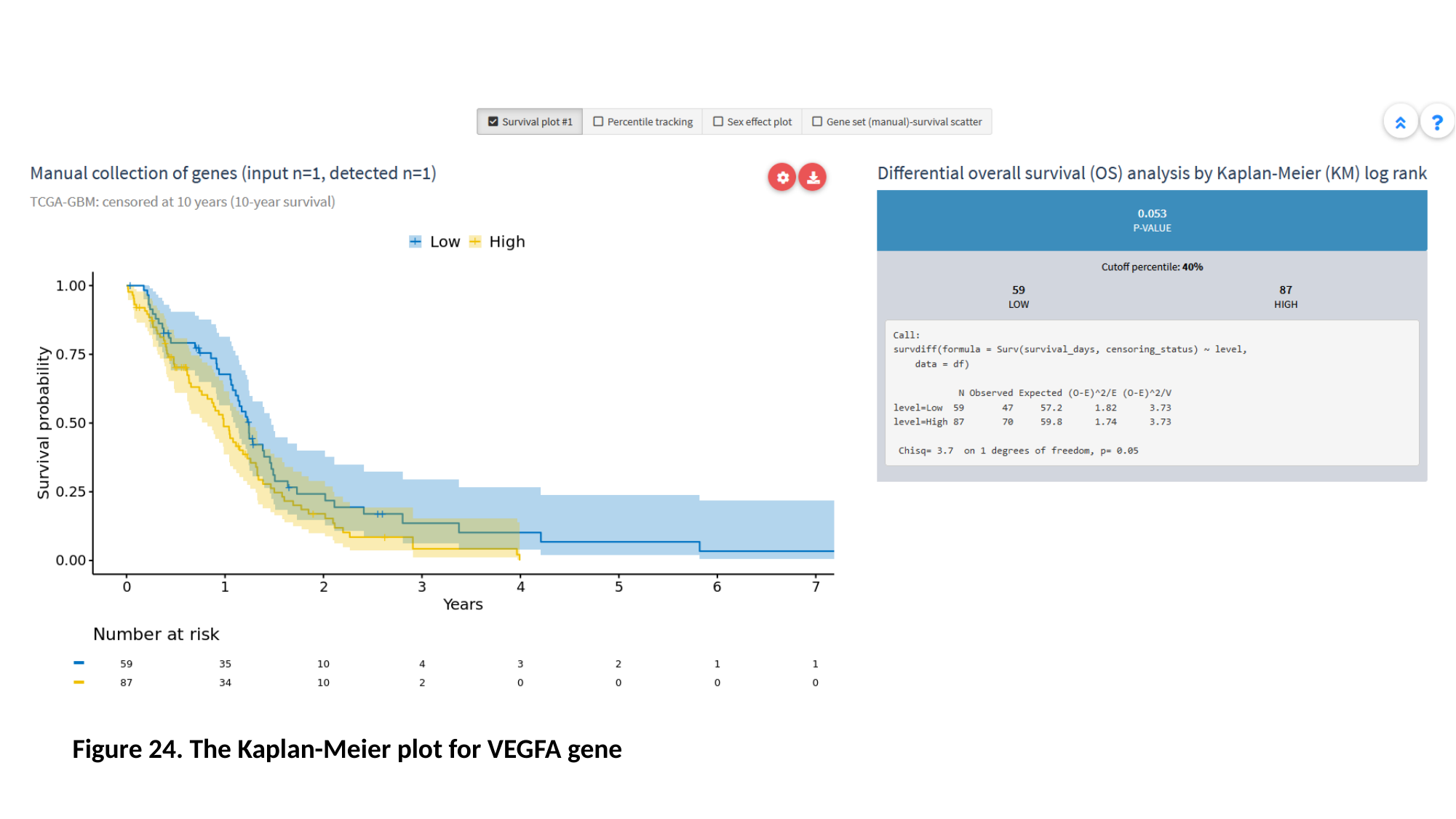

Figure 24. The Kaplan-Meier plot for VEGFA gene

### Slide 28
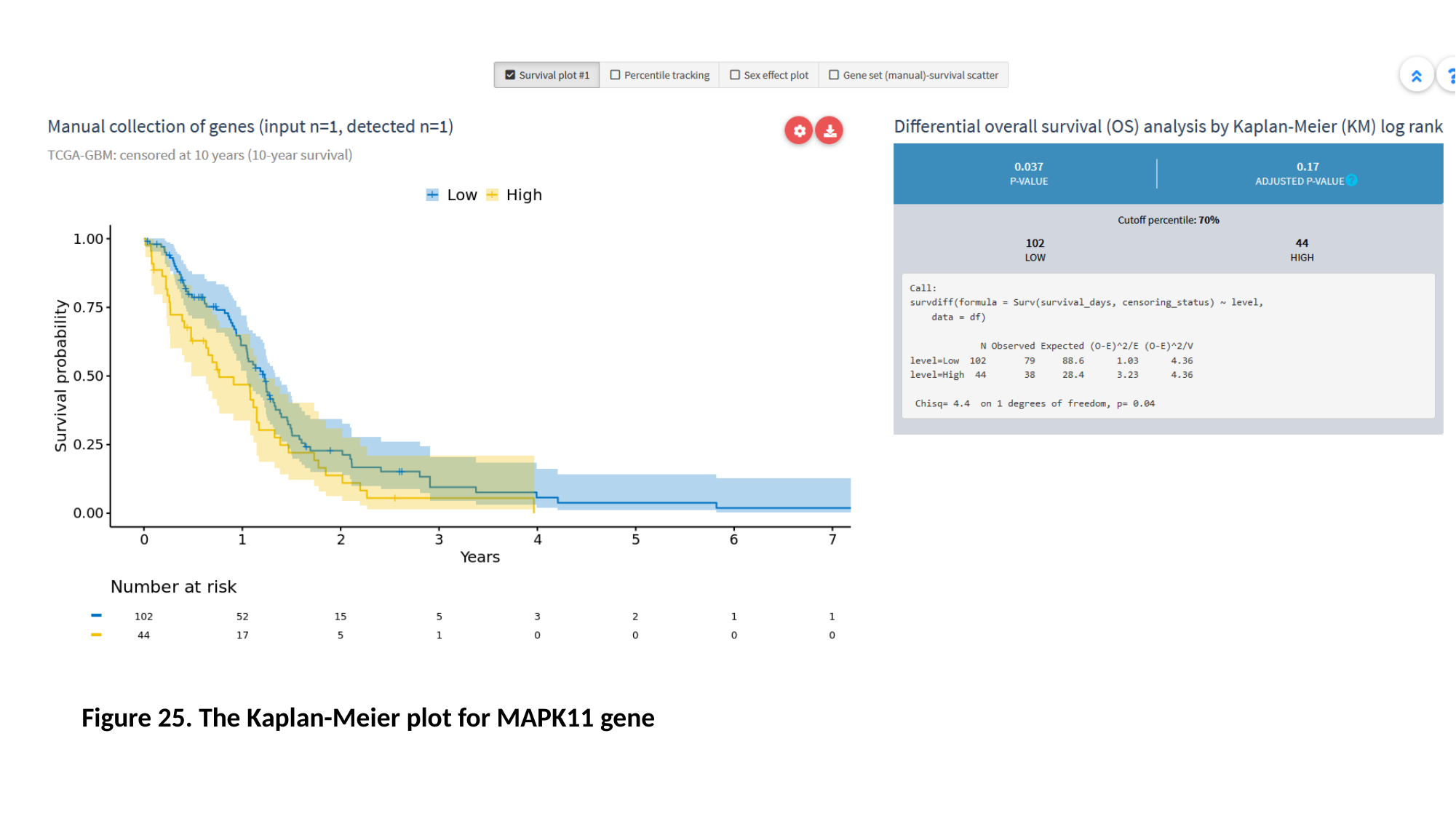

Figure 25. The Kaplan-Meier plot for MAPK11 gene

### Slide 29
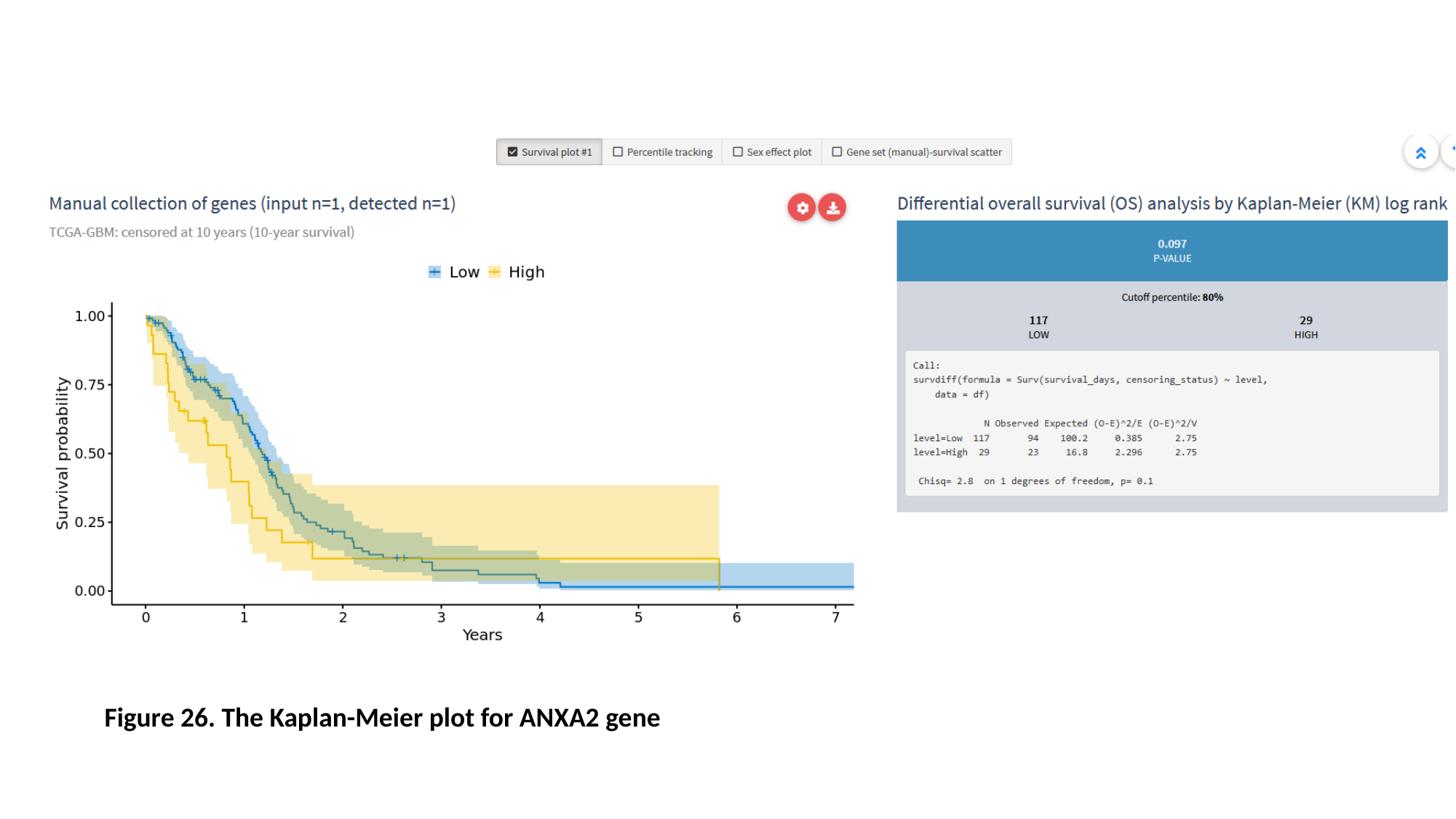

Figure 26. The Kaplan-Meier plot for ANXA2 gene

### Slide 30
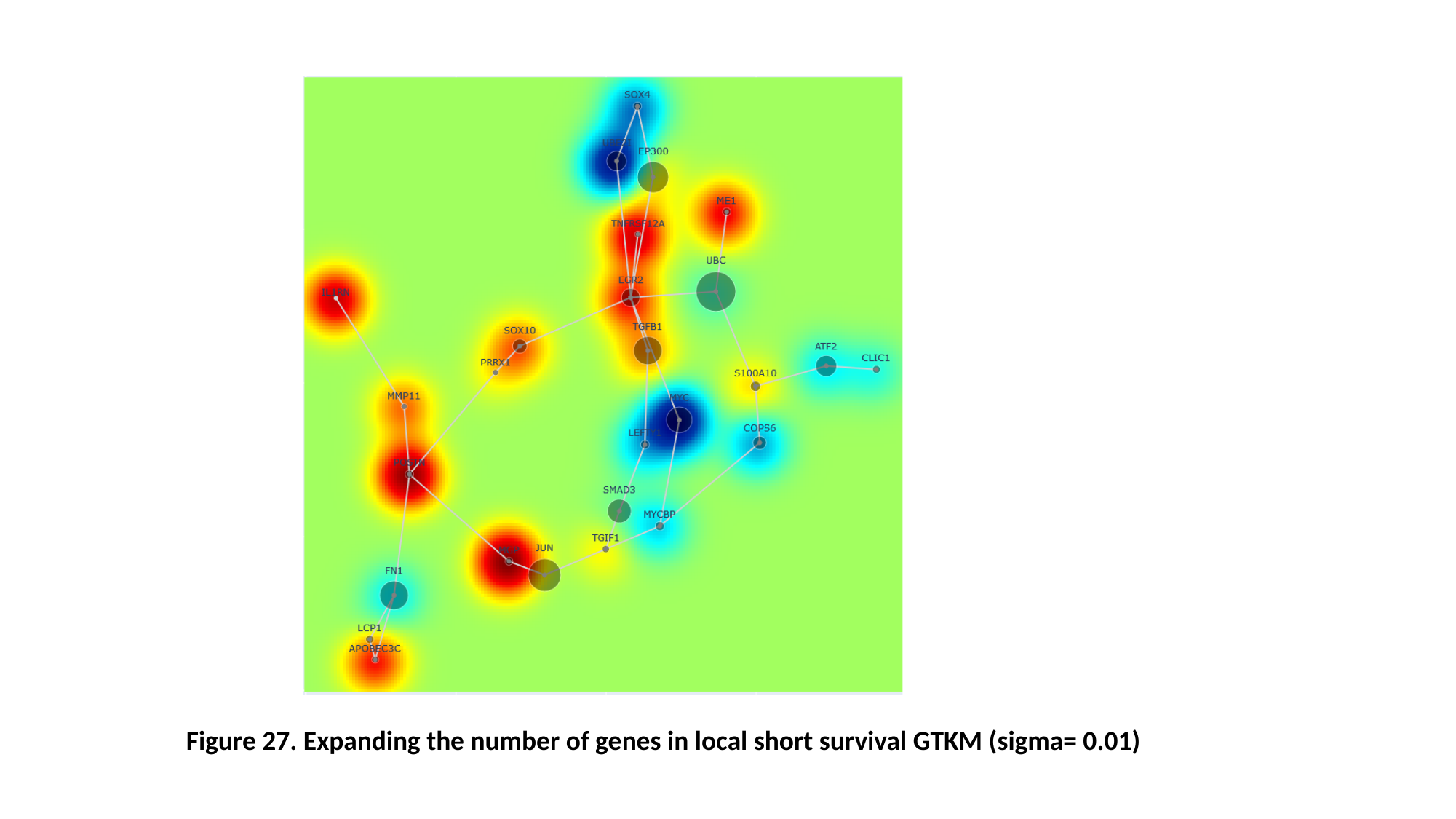

Figure 27. Expanding the number of genes in local short survival GTKM (sigma= 0.01)

### Slide 31
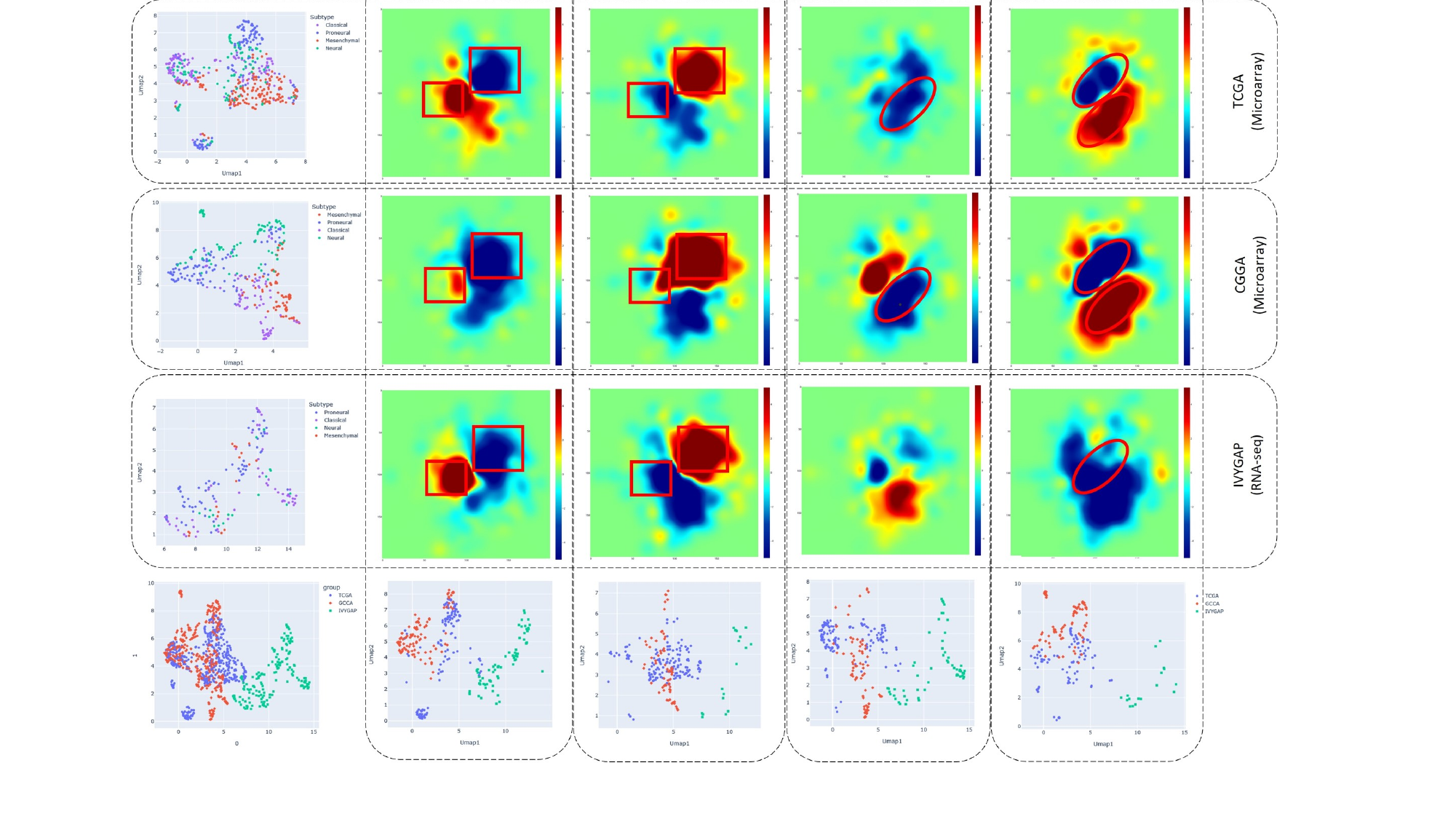

### Slide 32
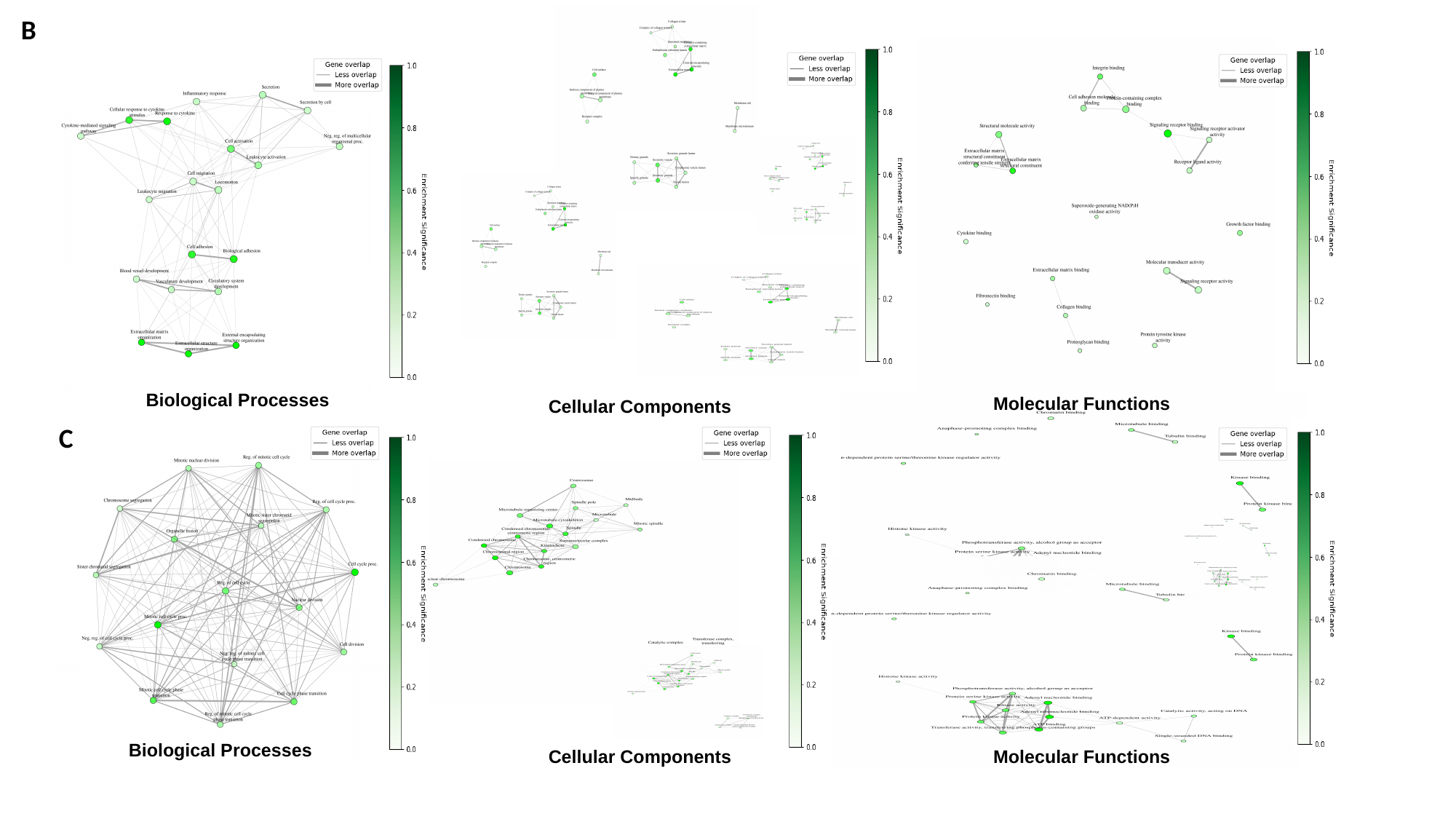

B
Biological Processes
Molecular Functions
Cellular Components
C
Biological Processes
Molecular Functions
Cellular Components

### Slide 33
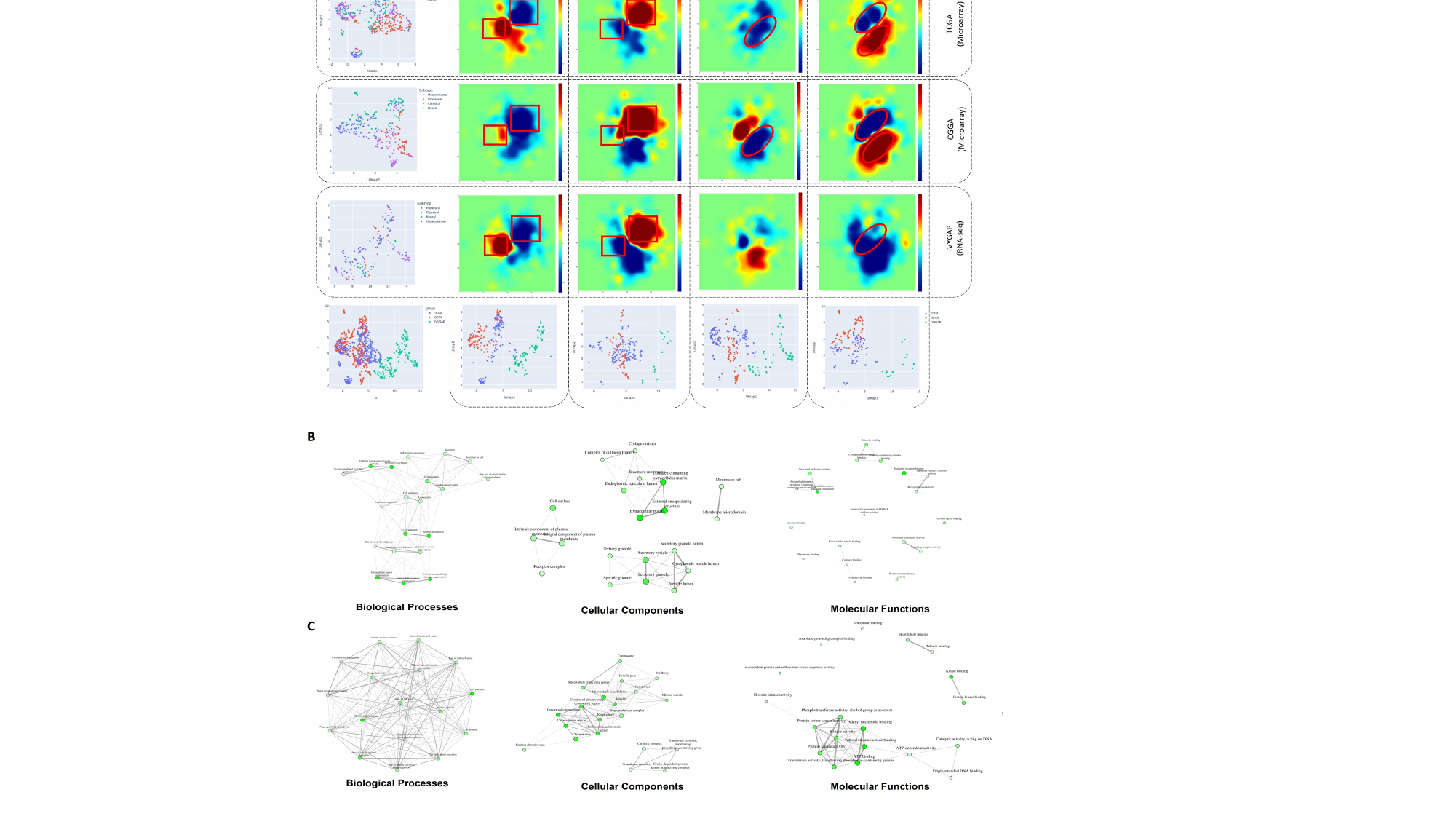

### Slide 34
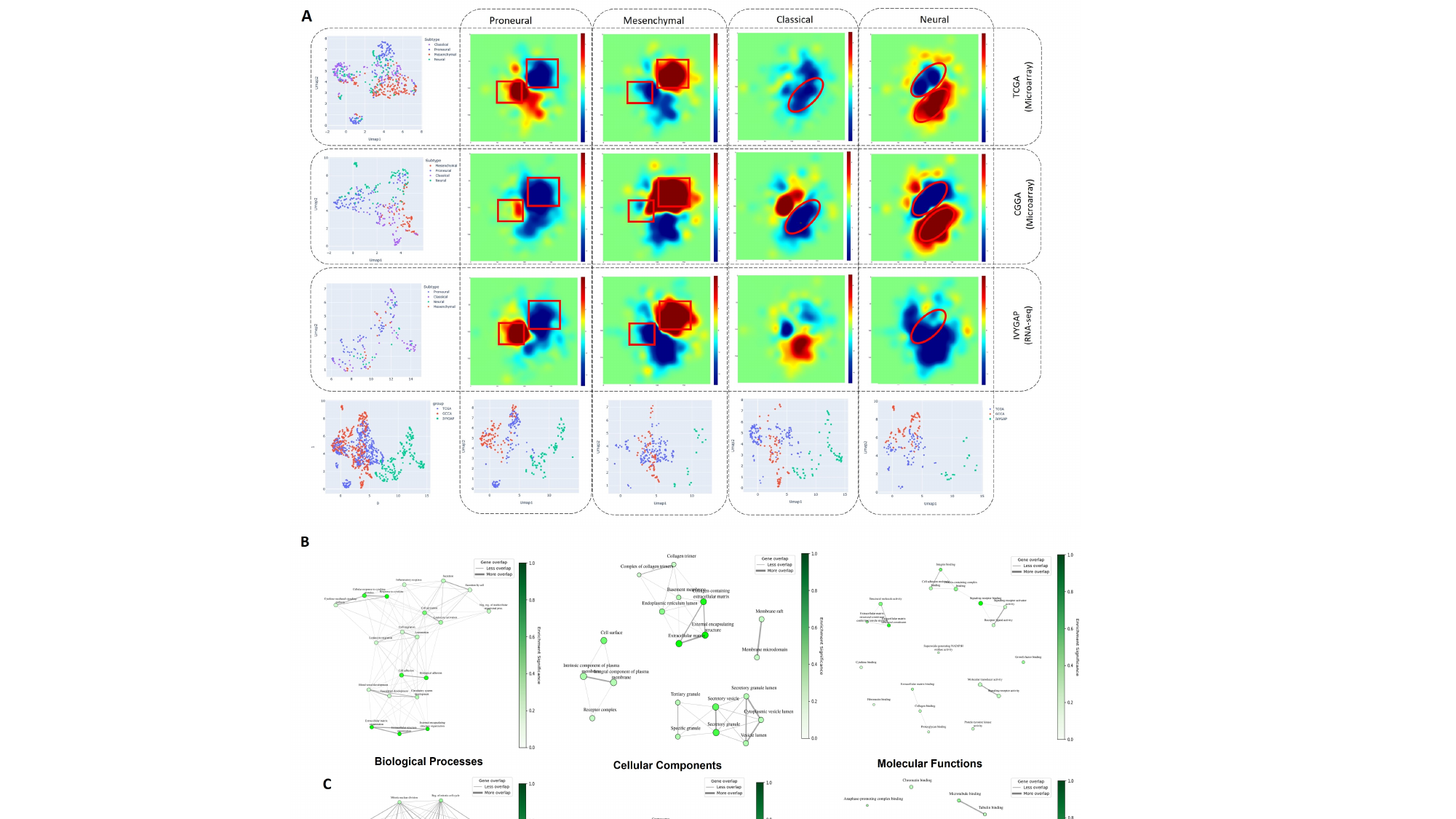
